# Human neutralizing antibodies to cold linear epitopes and to subdomain 1 of SARS-CoV-2

**DOI:** 10.1101/2022.11.24.515932

**Authors:** Filippo Bianchini, Virginia Crivelli, Morgan E. Abernathy, Concetta Guerra, Martin Palus, Jonathan Muri, Harold Marcotte, Antonio Piralla, Mattia Pedotti, Raoul De Gasparo, Luca Simonelli, Milos Matkovic, Chiara Toscano, Maira Biggiogero, Veronica Calvaruso, Pavel Svoboda, Tomás Cervantes Rincón, Tommaso Fava, Lucie Podešvová, Akanksha A. Shanbhag, Andrea Celoria, Jacopo Sgrignani, Michal Stefanik, Vaclav Hönig, Veronika Pranclova, Tereza Michalcikova, Jan Prochazka, Giuditta Guerrini, Dora Mehn, Annalisa Ciabattini, Hassan Abolhassani, David Jarrossay, Mariagrazia Uguccioni, Donata Medaglini, Qiang Pan-Hammarström, Luigi Calzolai, Daniel Fernandez, Fausto Baldanti, Alessandra Franzetti-Pellanda, Christian Garzoni, Radislav Sedlacek, Daniel Ruzek, Luca Varani, Andrea Cavalli, Christopher O. Barnes, Davide F. Robbiani

## Abstract

Emergence of SARS-CoV-2 variants diminishes the efficacy of vaccines and antiviral monoclonal antibodies. Continued development of immunotherapies and vaccine immunogens resilient to viral evolution is therefore necessary. Using coldspot-guided antibody discovery, a screening approach that focuses on portions of the virus spike that are both functionally relevant and averse to change, we identified human neutralizing antibodies to highly conserved viral epitopes. Antibody fp.006 binds the fusion peptide and cross-reacts against coronaviruses of the four *genera*, including the nine human coronaviruses, through recognition of a conserved motif that includes the S2’ site of proteolytic cleavage. Antibody hr2.016 targets the stem helix and neutralizes SARS-CoV-2 variants. Antibody sd1.040 binds to subdomain 1, synergizes with antibody rbd.042 for neutralization and, like fp.006 and hr2.016, protects mice when present as bispecific antibody. Thus, coldspot-guided antibody discovery reveals donor-derived neutralizing antibodies that are cross-reactive with *Orthocoronavirinae*, including SARS-CoV-2 variants.

**One sentence summary:** Broadly cross-reactive antibodies that protect from SARS-CoV-2 variants are revealed by virus coldspot-driven discovery.

---

The coronavirus (CoV) spike protein (S) is a trimeric glycoprotein of S1-S2 heterodimers that mediates binding to target cells and membrane fusion (*1–3*). Most severe acute respiratory syndrome coronavirus 2 (SARS-CoV-2) neutralizing antibodies that have been described to date target the receptor binding and N-terminal domains of S (RBD and NTD) (*4–6*). However, mutations in the viral genome, such as those found in SARS-CoV-2 variants of concern (VOC), cause amino acid (aa) changes in the RBD and NTD that diminish or abrogate the effectiveness of vaccines and antiviral monoclonal antibodies currently in the clinic (*7–10*). Thus, innovative approaches are needed to identify countermeasures that remain effective in spite of SARS-CoV-2 viral evolution and have the potential to combat the growing number of coronaviruses that cause infection in humans (*11–13*).

## Results

We hypothesized that some regions of S may be under selective pressure to maintain their aa sequence unchanged because it is essential for their function or to maintain proper quaternary structure. To determine if such regions exist, we analyzed 10,480,461 SARS-CoV-2 sequences from GISAID (*14*). We identified 15 regions with infrequent aa changes (coldspots, defined as >17 consecutive aa with frequency of substitutions <0.1%): one coldspot includes the S2’ cleavage site and a portion of the fusion peptide (FP, aa 814–838), which is substrate of the TMPRSS2 and Cathepsin proteases; a second one is at the stem helix that precedes the heptad repeat 2 region (HR2, aa 1142–1161); and three coldspots span sequences at the discontinuously encoded subdomain 1 (SD1; see table S1). Both FP and HR2 coldspots are devoid of aa changes in SARS-CoV-2 VOC, while changes are rare in SD1 (Fig. 1, A and B and fig. S1A).

**Fig. 1.**
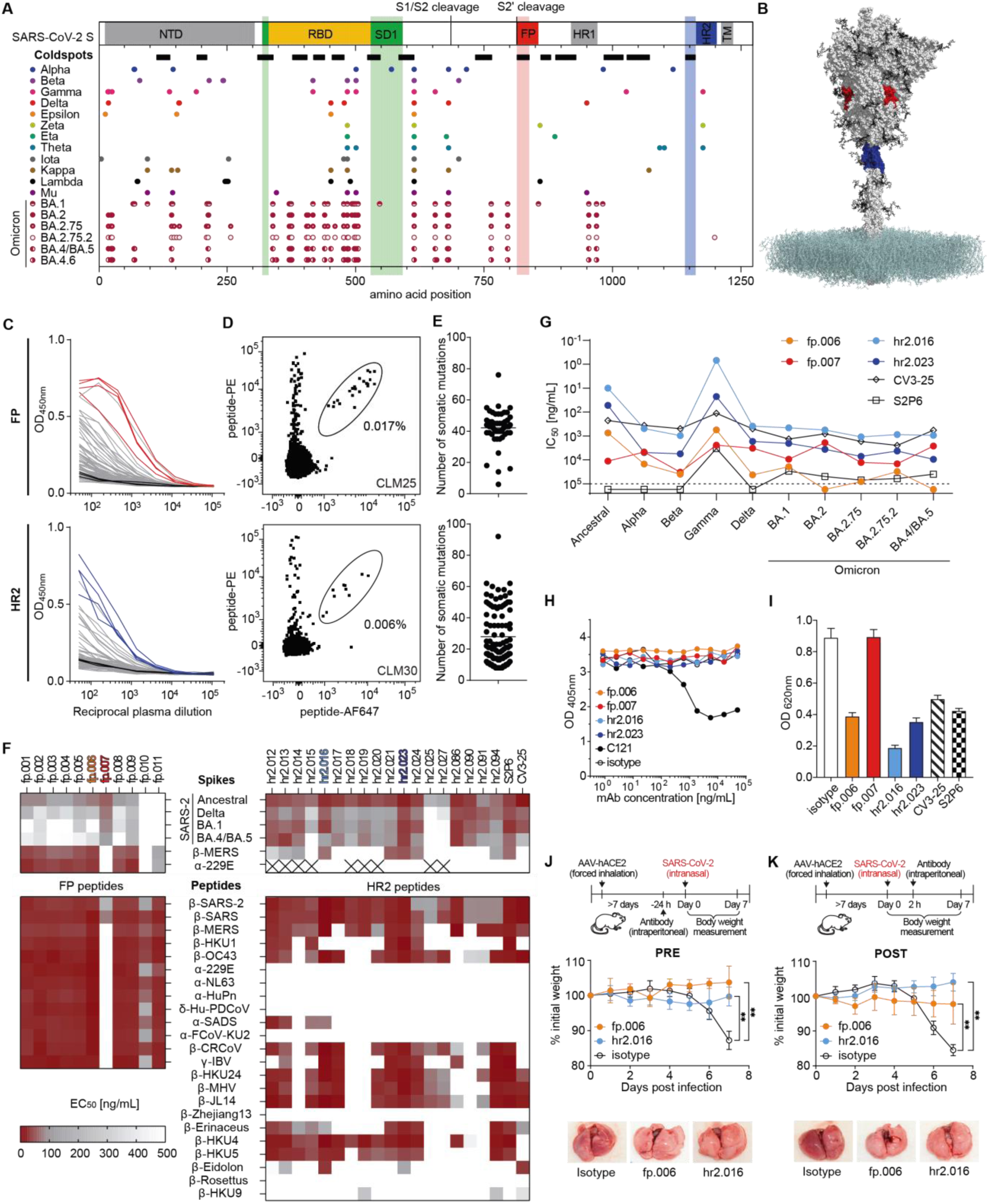
Discovery of virus-neutralizing coldspot antibodies. **(A)** On top, cartoon diagram of the SARS-CoV-2 spike with highlighted the coldspot areas at the fusion peptide (FP, red), heptad repeat 2 region (HR2, blue), and subdomain 1 (SD1, green). The thick horizontal lines indicate the location of all coldspots (see also fig. S1A). At the bottom, amino acid changes in SARS-CoV-2 variants. Each circle represents a single aa substitution over ancestral virus. **(B)** Structure of the SARS-CoV-2 spike; FP (aa 814-838) and HR2 (aa 1142-1161) coldspots are in red and blue, respectively (PDB: 6XM4). **(C)** Graph shows ELISA measurements of convalescent plasma IgG reactivity to FP (top) or HR2 (bottom) peptides. Optical density units at 450 nm (OD, Y axis) and reciprocal plasma dilutions (X axis). Non-infected controls in black; samples selected for cell sorting by flow cytometry are in red or blue. Two independent experiments. **(D)** Representative flow cytometry plots of B cells binding to fluorescently labeled FP (top) or HR2 (bottom) peptides. Numbers indicate percentage of double-positive cells in the gate. **(E)** Number of heavy and light chain V gene somatic mutations of antibodies to the FP (top) or HR2 (bottom) peptides. **(F)** Heatmaps with ELISAs EC_50_ values of monoclonal antibodies binding to the S of CoVs (top) and to the FP and HR2 peptides (bottom) corresponding to the CoV species, whose genus is indicated by Greek letters. The monoclonal antibodies to the HR2 region S2P6 (*16*) and CV3-25 (*17*) were assayed alongside for comparison. Cross indicates not tested. Two experiments. **(G)** Graph with IC_50_ values of monoclonal antibodies neutralizing pseudoviruses corresponding to the indicated VOC. Two experiments. **(H)** ACE2 binding to ancestral S in ELISA in the presence of select FP and HR2 antibodies. Dotted line represents the limit of detection. Two experiments. **(I)** Inhibition of cell fusion by FP and HR2 antibodies. Two experiments. **(J** and **K)** fp.006 and hr2.016 antibodies protect in vivo. Top, diagram of the experiment’s timeline. Middle, mouse weight over time after challenge with ancestral SARS-CoV-2 of mice treated with antibodies either 24 hours before (**(J)**; n=6 per group, p=0.0022 for both fp.006 and hr2.016 versus isotype at day 7), or 2 hours after (**(K)**; n=5 per group, p=0.0079 for both fp.006 and hr2.016 versus isotype at day 7) the infection. Mann-Whitney U test, standard deviation is shown. At the bottom, representative lung images at day 7.

To determine whether antibodies to the FP and HR2 coldspots occur naturally, we evaluated plasma samples from a COVID-19 convalescent cohort by enzyme-linked immunosorbent assay (ELISA; n=67; see Methods). High levels of IgG antibodies binding to peptides at the FP and HR2 coldspots were found in convalescent individuals (Fig. 1C). In comparison, IgG levels were low to undetectable in samples from uninfected controls, in pre-pandemic samples obtained after documented common cold CoV infection, and in most COVID-19 vaccinated individuals, except for some who received inactivated virus-based vaccines (fig. S1B). To examine the molecular features of coldspot antibodies, we used flow cytometry to isolate B cells specific for FP and HR2 peptides from those individuals with high antibody levels in plasma (Fig. 1D, fig. S1, C and D; see Methods). We obtained 55 (FP) and 100 (HR2) paired IgG heavy and light chain antibody sequences, some of which clustered in expanded clones of related antibodies (fig. S1E). The average number of V gene somatic nucleotide mutations was high: 42 for FP (range: 6-76) and 28 for HR2 antibodies (range: 8-92; Fig. 1E).

Twenty-nine monoclonal antibodies, including at least one representative for each of the expanded clones, were recombinantly expressed and tested in ELISA (table S1). Ten out of 11 FP antibodies bound to the peptide with half-maximal effective concentrations (EC_50_) between 25-119 ng/mL. The EC_50_ values of the same antibodies to S were on average 4.8-fold higher, except for antibody fp.007, where the observed EC_50_ declined from 177 to 58 ng/mL (Fig. 1F and fig. S2). Similarly, all 18 HR2 antibodies bound to the peptide with EC_50_ values between 7-117 ng/mL, and with an average of 1.3-fold higher EC_50_ to S. Several of the antibodies were broadly cross-reactive since they bound to representative S trimers of SARS-CoV-2 VOC, MERS and HCoV-229E, as well as to FP and HR2 peptides corresponding to other CoVs. Noteworthy, most FP antibodies recognized CoVs of the four *genera* (alpha to delta, including all 9 CoVs associated with human disease), and some HR2 antibodies cross-reacted not only with beta-, but also with alpha- and gammacoronaviruses (Fig. 1F, figs. S2 and S3; table S2) (*11, 15, 16*).

**Fig. 2.**
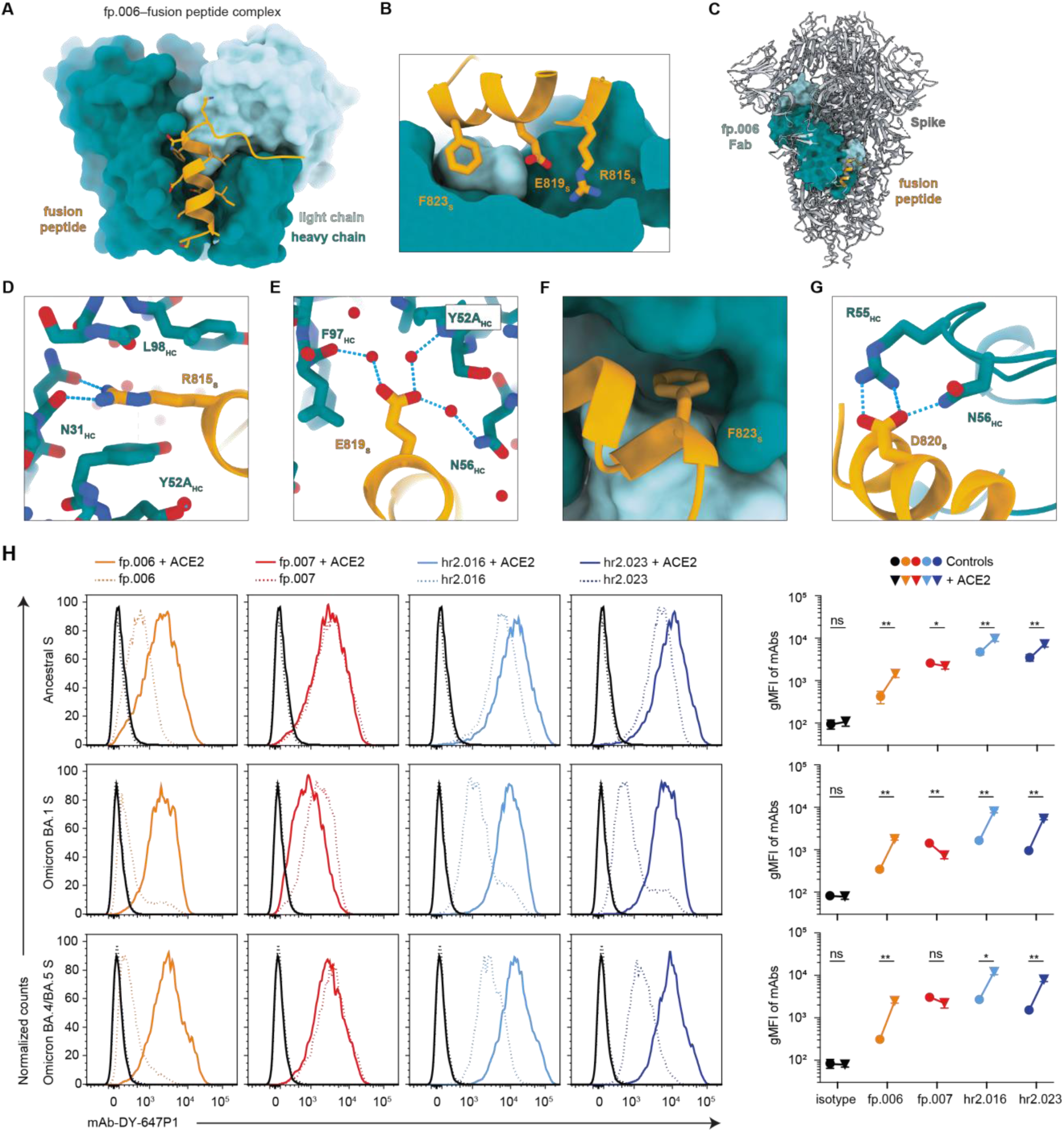
SARS-CoV-2 FP recognition by fp.006. **(A)** Overview of the complex structure of fp.006 Fab (surface representation; heavy chain in teal, light chain in light teal) bound to the SARS-CoV-2 FP (orange cartoon) with interacting side chains represented as sticks. **(B)** Visualization of FP residues F_823_, E_819_, and R_815_ resting in a deep groove formed at the antibody paratope, with coloring as in (**A**). **(C)** Overlay of the fp.006-FP crystal structure with a cryo-EM structure of the SARS-CoV-2 prefusion S trimer (PDB: 6VXX). Models were aligned on Cα atoms of FP residues 818-822 (helical in both structures) with a root mean square deviation of 0.97 Å. **(D)** Residue-level interactions between FP residue R_815_ and the antibody heavy chain include hydrogen bond formation with N_31_ and a cation-π interaction with Y_52A_. **(E)** Water-mediated interactions between FP residue E_819_ and heavy chain residues Y_52A_, N_56_, and F_97_. Water molecules are shown as red spheres. **(F)** van der Waals contacts between FP residue F_823_ (orange stick) and residues that comprise a groove at the heavy and light chain interface (teal surfaces). **(G)** Interactions between FP residue D_820_ and fp.006 CDRH2 residues include a salt bridge with R_55_ and additional hydrogen bond formation with N_56_. Hydrogen bonds, salt bridges, and cation-π interactions are shown as dashed blue lines. **(H)** Flow cytometry detection of anti-FP and anti-HR2 antibody binding to SARS-CoV-2 S expressed on 293T cells. Left, representative FACS plots (pre-gated on live-singlets-GFP_+_ cells). Black lines indicate isotype control in the presence (continuous line) or absence (dotted line) of soluble ACE2. Right, quantification of the geometric mean fluorescent intensity (gMFI; n=4). Two-tailed paired t-test: *p<0.05, **p<0.01, ***p<0.001 and ****p<0.0001; standard deviation is shown.

**Fig. 3.**
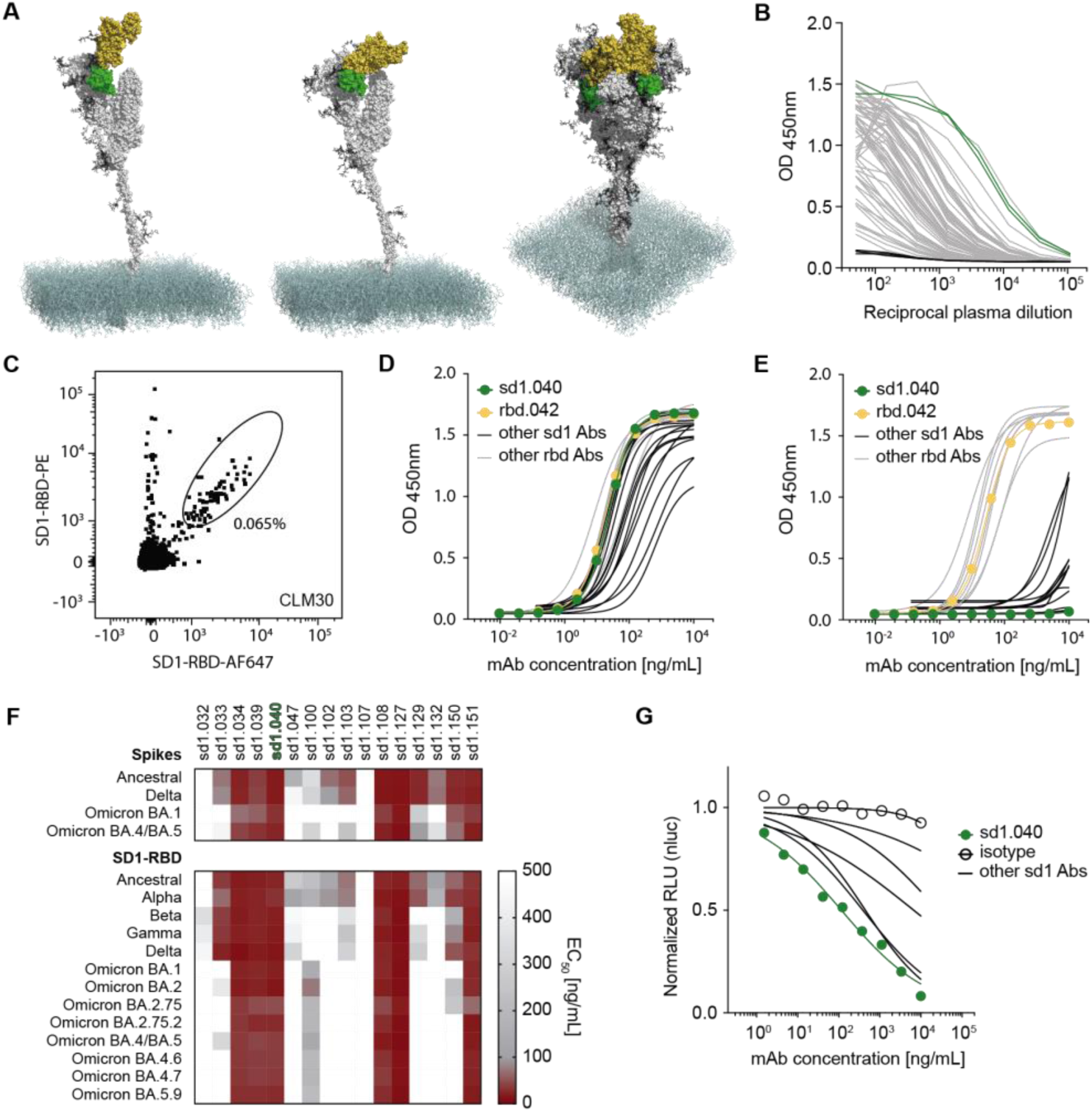
Discovery of broadly cross-reactive antibodies to SD1 and RBD. **(A)** Structure of the SARS-CoV-2 S. S protomer with RBD up (left) or down (middle) and S trimer with two down and one up (right; PDB: 6XM4). SD1 and RBD are in green and yellow, respectively. **(B)** Graph shows ELISAs measuring plasma IgG reactivity to SD1-RBD. Negative controls in black; samples selected for sorting in green. Mean of two independent experiments. 82.1% of the plasma samples were positive (4SD higher than the average AUC of the controls) **(C)** Representative flow cytometry plot of B cells binding to fluorescently labeled SD1-RBD. Percentage refers to gated cells. **(D and E)** ELISAs measuring the reactivity of monoclonal antibodies to SD1-RBD (**D**) and to RBD (**E**). Mean of two independent experiments. **(F)** Heatmaps with the binding (EC_50_) of SD1 monoclonal antibodies to S (top) or SD1-RBD (bottom) proteins corresponding to SARS-CoV-2 VOC. Two experiments. **(G)** Graph shows normalized relative luminescence values in cell lysates of 293T_ACE2_ cells after infection with ancestral SARS-CoV-2 pseudovirus in the presence of increasing concentrations of broadly crossreactive SD1 monoclonal antibodies. At least two independent experiments.

To evaluate the antibodies’ ability to neutralize SARS-CoV-2, we used a previously established SARS-CoV-2 pseudovirus assay (*4*). The most potent FP antibody (fp.006) displayed a half-maximal inhibitory concentration (IC_50_) of 737 ng/mL, while the best HR2 neutralizer (hr2.016) had an IC_50_ of 10 ng/mL, which was lower than previously reported antibodies to this region that were tested alongside (CV3-25 and S2P6; Fig. 1G; S4A; table S2) (*16, 17*). Select anti-FP and anti-HR2 antibodies blocked infection regardless of TMPRSS2-expression by target cells and, consistent with the view that they antagonize post-attachment events, they did not interfere with ACE2 binding to S in ELISA but inhibited cell fusion (Fig. 1, H and I, fig. S4, A and B). As expected, based on the absence of aa changes at coldspot regions, some FP and HR2 antibodies were effective against pseudoviruses corresponding to SARS-CoV-2 VOC, against ancestral and Omicron SARS-CoV-2 *in vitro*, and protected mice challenged with ancestral virus when administered either as pre- or post-exposure prophylaxis (Fig. 1, G, J and K; fig. S4, C and D; table S2). Thus, natural antibodies exist that protect against SARS-CoV-2 by binding to highly conserved linear epitopes at functional regions of S.

**Fig. 4.**
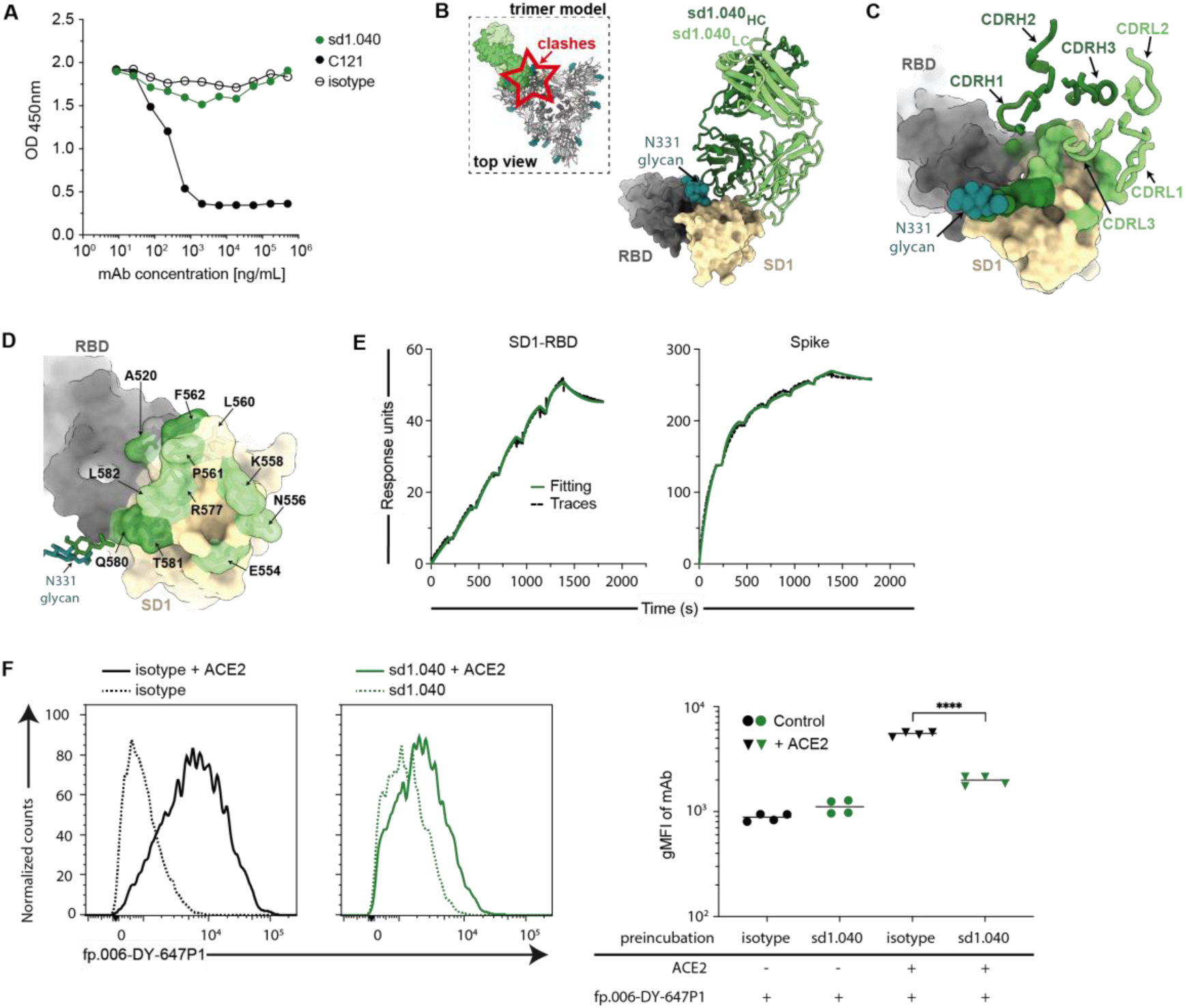
Cryo-EM structure of sd1.040 in complex with SARS-CoV-2 S. **(A)** ACE2 binding to ancestral S in ELISA in the presence of sd1.040 or C121 control antibody. Representative of two experiments. **(B)** Structure of the sd1.040-S complex. Spike SD1 and RBD regions are shown as surface representation and colored wheat and gray, respectively. The sd1.040 Fab heavy chain (dark green) and light chain (light green) are shown as cartoon. The S N331-glycan that interacts with the sd1.040 Fab is shown as teal spheres. Inset: sd1.040 binding orientation on trimeric S shows clashes. **(C and D)** Surface rendering of sd1.040 epitope is highlighted on the SD1 and RBD surfaces, with sd1.040 CDR loops shown (ribbon). The majority of sd1.040 contacts are mediated by CDRH2, CDRL1 and CDRL3 loops. **(E)** Surface plasmon resonance (SPR) experiment showing the binding of sd1.040 Fab to ancestral SD1-RBD or S. **(F)** Antibody sd1.040 prevents ACE2-induced rearrangements. Flow cytometry detection of fp.006 binding to ancestral SARS-CoV-2 S expressed on 293T cells. Left, representative FACS plots. Black lines indicate isotype control in the presence (continuous line) or absence (dotted line) of soluble ACE2. Right, quantification of the geometric mean fluorescent intensity (gMFI; n=4). Two-tailed paired t-test: *p<0.05, **p<0.01, ***p<0.001 and ****p<0.0001.

To gain insight into the mechanism of broad recognition by fp.006, we obtained a 2.0 Å resolution crystal structure of its Fab in complex with the _812_PSKRSFIEDLLFNKVTLADA_831_ FP peptide (Fig. 2, table S3). In the bound structure, FP residues 813-825 adopt an α-helical conformation, extending an amphipathic helix observed in prefusion S trimer structures. The helical peptide sits within a groove that is formed by fp.006 complementarity-determining region (CDR) 3 loops, which mediate most epitope contacts (Fig. 2A). Additional contacts with heavy chain CDR1 and CDR2 loops result in a total buried surface area (BSA) of 1,466 Å^2^ (686 Å^2^ paratope BSA + 780 Å^2^ epitope BSA) and are consistent with binding orientations of similarly described anti-FP antibodies (*18–20*) (fig. S5, A to G). Of the 13 antigenically distinct CoV FPs tested here, the majority of epitope residues contacted by fp.006 are highly conserved, explaining fp.006’s breadth of binding (Fig. 1F; fig. S2B and S5B). In particular, three FP residues (R_815_, E_819_, and F_823_) are completely buried in the Fab groove and make extensive hydrogen bond and hydrophobic interactions (Fig. 2, B to G). As a result, one face of the FP amphipathic helix comprises two polar residues that contact a polar patch on the edge of the Fab trough formed by CDRH2, and the opposite hydrophobic face engages hydrophobic residues in the CDRH3 loop (fig. S5, D and E). Notably, residue R_815_, which is critical for TMPRSS2 and Cathepsin cleavage (*3, 21, 22*), forms hydrogen bonds with the Fab CDRH1 loop and a cation-π interaction with Y_52A_ in CDRH2 (Fig. 2D). Given the importance of R_815_ in protease cleavage, fp.006-mediated neutralization likely includes steric hinderance of TMPRSS2 and Cathepsin binding and further processing of the S trimer for productive fusion. Interestingly, superposition of the Fab-FP complex crystal structure with a prefusion S trimer structure revealed an approach angle incompatible with Fab binding, which explains the weak binding observed for FP antibodies to prefusion S trimers (Fig. 1F and Fig. 2C). Thus, cleavage by cellular proteases at the S2’ site and antibody recognition of this partially cryptic epitope likely involves transient conformational changes that are necessary to expose the FP epitope.

**Fig. 5.**
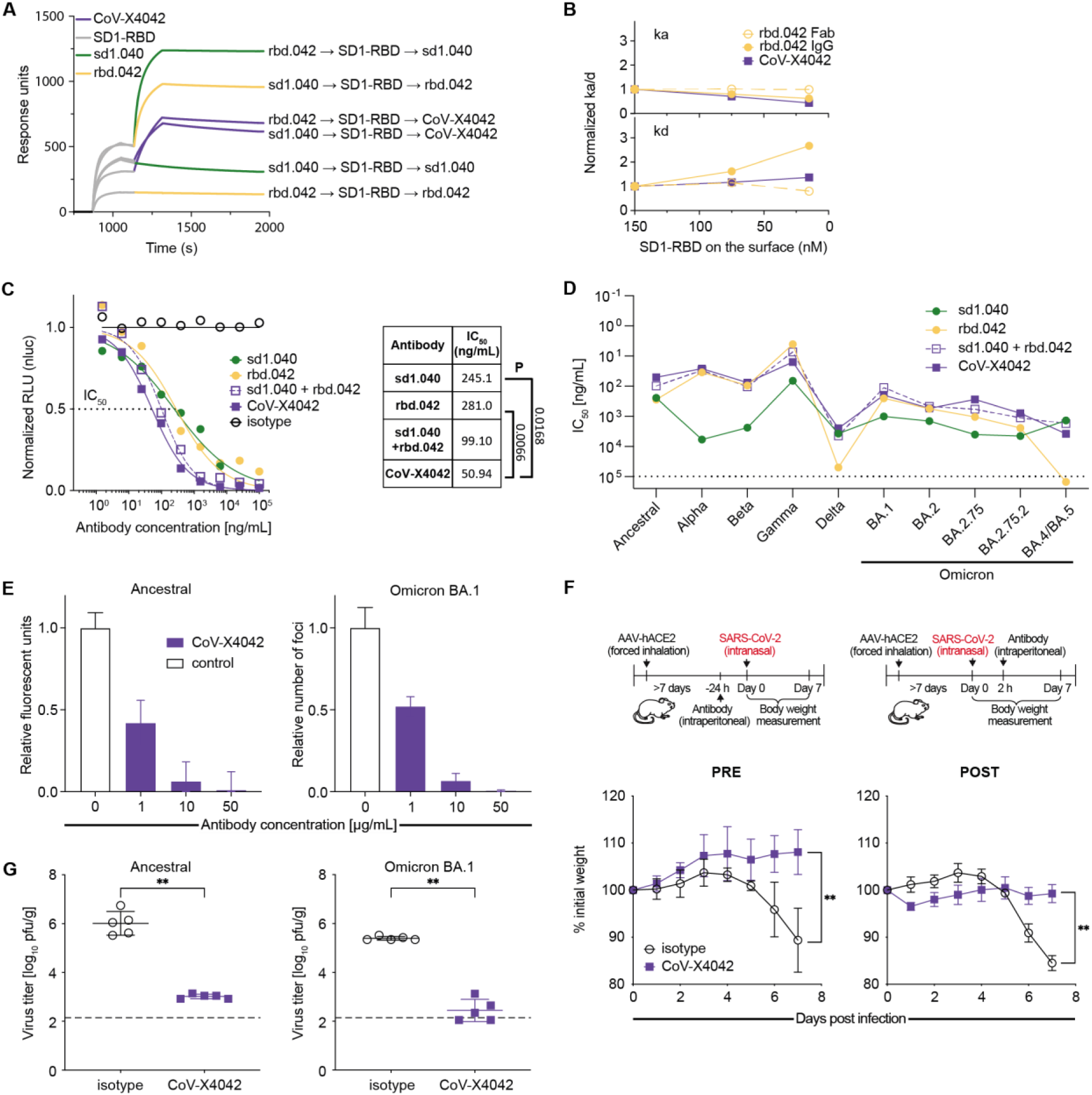
In vitro neutralization and mouse protection by the bispecific antibody CoV-X4042. **(A)** SPR assay of the sequential binding of immobilized antibodies to SD1-RBD protein followed by either sd1.040, rbd.042 or CoV-X4042. **(B)** SPR analysis showing that both arms of CoV-X4042 bind simultaneously to the same SD1-RBD molecule, since avidity is retained at decreasing SD1-RBD concentrations. Increasing normalized kd values indicate loss of avidity. Solid lines, IgG; dotted lines, Fab (see also fig. S8A). **(C)** Normalized relative luminescence values in cell lysates of 293T_ACE2_ cells after infection with ancestral SARS-CoV-2 pseudovirus in the presence of increasing concentrations of CoV-X4042 or its parental monoclonal antibodies individually or as a cocktail. Isotype control in black. On the right: mean IC_50_ values and significance (P) when parental antibodies are compared to CoV-X4042 (n=4; Welch’s t-test, two-tailed). **(D)** Graph with IC_50_ values of bispecific and parental monoclonal antibodies neutralizing pseudoviruses corresponding to the indicated VOC. Mean of two independent experiments. **(E)** In vitro neutralization of SARS-CoV-2 by CoV-X4042. **(F)** CoV-X4042 protects in vivo. Top, diagram of the experiment’s timeline. Bottom, mouse weight over time after challenge, with ancestral SARS-CoV-2, of mice treated with antibodies either 24 hours before (PRE; n=5 per group, p=0.0079), or 2 hours after (POST; n=5 per group, p=0.0079; day 7) the infection. Mann-Whitney U test, standard deviation is shown. **(G)** CoV-X4042 reduces viral titers in the lungs. Mice were treated with antibodies 24 hours before infection and virus titers evaluated on day 3 (n=5 per group p=0.0079 with both ancestral and Omicron BA.1; Mann-Whitney U test, standard deviation is shown).

In agreement with this view, ACE2 engagement of cell surface-expressed S, which is known to alter S conformation, increased fp.006 binding by 5.8-fold in a flow cytometry assay, and the addition of soluble ACE2 synergized with fp.006 for neutralization (Fig. 2H and fig. S5, H and I). Interestingly, and consistent with the ELISA and neutralization data (Fig. 1, F and G), binding of fp.006 to S was weaker with Omicron by this assay, but ACE2 attachment improved it to similar levels as with ancestral (5.3-fold for BA.1 and 8.2-fold for BA.4/BA.5; Fig. 2H). Therefore, ACE2 can induce conformational changes of S that expose the highly conserved FP epitope and favor neutralization. ACE2 attachment also increased the binding of hr2.016 and hr2.023 to both ancestral (2.3- and 2.4-fold) and Omicron S (4.9-fold and 5.8-fold for BA.1, 4.6-fold and 5.3-fold for BA.4/BA.5, respectively; Fig. 2H), but did not improve the binding of fp.007, a neutralizing FP-antibody that displays a different pattern of cross-reactivity than fp.006 (Fig. 1F and 2H). Therefore, optimal FP recognition by neutralizing fp.007-like antibodies does not require ACE2-induced conformational changes.

The subdomain 1 (SD1) of SARS-CoV-2 S is adjacent to the RBD and its sequence is conserved across SARS-CoV-2 variants, except for substitutions A570D in Alpha (B.1.1.7) and T547K in Omicron BA.1 (B.1.1.529; Fig. 1, A and B and 3A). To identify antibodies targeting SD1, we designed a flow cytometry-based strategy that combines negative selection of B cells binding to RBD (aa 331-529) with positive selection of those binding to SD1-RBD (aa 319-591; fig. S1C; see Methods). Peripheral blood mononuclear cells were obtained from individuals with high plasma IgG reactivity to SD1-RBD (Fig. 3B and fig. S6, A and B) and B cells enriched for binding to SD1 were sorted as single cells for antibody gene sequencing (Fig. 3C, fig. S6C).

Twenty-five monoclonal antibodies were cloned and produced. All 25 bound to SD1-RBD in ELISA, and 16 were SD1-specific (Fig. 3, D and E and table S1). Six of the SD1 antibodies cross-reacted with SD1-RBD proteins corresponding to all twelve CoV VOC with EC_50_ values of 66.25 ng/mL or lower, while only two RBD antibodies cross-reacted with all variants’ RBD at low EC_50_ (Fig. 3F; fig. S6, D to F, and table S2). Select antibodies also bound effectively to the common SD1 variants T572I and E583D (fig. S6G). In pseudovirus-based neutralization assays, the best broadly cross-reactive SD1 antibody was sd1.040 (IC_50_=245 ng/mL; Fig. 3G, and table S2). This demonstrates that naturally occurring antibodies can neutralize SARS-CoV-2 by binding to SD1.

The mechanism of neutralization by sd1.040 does not involve inhibition of receptor binding, since sd1.040, unlike C121 (*4*), failed to prevent ACE2 binding to S in ELISA (Fig. 4A). To gain insight into the neutralization mechanism, we formed a complex between sd1.040 Fabs bound to a prefusion S-2P trimer and used cryo-electron microscopy (cryo-EM) for structure determination. Interestingly, no intact Fab-S trimer structures were observed within the dataset (fig. S7 A to D). Instead, our 3.7 Å cryo-EM structure revealed sd1.040 Fabs in complex with S1 protomers, recognizing an epitope comprising SD1 residues 554-562, 577-581, and RBD residues 520-524 (Fig. 4, B to D, fig. S7 A to D, and table S4). Superposition of the cryo-EM sd1.040-S protomer complex on published prefusion S structures, indiscriminate of RBD conformation, revealed minor clashes with the N-terminal domain of the adjacent protomer similar to antibody P008_60 (*23*) which contrasts the recently described SD1-specific murine antibody S3H3 (*24*) (Fig. 4B, inset, and fig. S7, E and F). It is likely that sd1.040 binding to S requires minor local rearrangement of the NTD. Indeed, docking and Molecular Dynamics computational simulations suggest the presence of quaternary interactions between the antibody, SD1 and RBD on one protomer, and NTD on another, which require minor rearrangement of the NTD hinging around residues 295-300 towards the end of the NTD (fig. S7, G and H). Such rearrangement does not result in clashes with other S regions. The CDRH1 and CDRH3, which do not interact with RBD/SD1 in the cryo-EM protomer structure, are shown to directly contact the NTD in the simulations. These quaternary interactions are likely to stabilize the complex, which by surface plasmon resonance (SPR) displays ∼20 times stronger binding affinity for S (*K*_D_ 0.2nM) than for SD1-RBD (*K*_D_ 4.2nM; Fig. 4E). To test the hypothesis that the mechanism of neutralization by sd1.040 is through the inhibition of S molecular rearrangements, we measured fp.006 binding to S by flow cytometry. ACE2 attachment in this assay improves fp.006 binding to S, which is however blocked by treatment with sd1.040 (Fig. 4F). Thus sd1.040 interferes with conformational changes downstream of ACE2 attachment.

Based on the breadth of binding exhibited by antibodies sd1.040 and rbd.042 against SARS-CoV-2 VOC, their potency of neutralization, and preliminary experiments that indicated synergistic effects from combining them (Fig. 3, F and G and fig. S6, F and H), we produced a bispecific antibody that includes the moieties of both antibodies, named CoV-X4042. CoV-X4042 has a natural IgG format including the entire

Fc region and complementary modifications in the Fc and CH1/CL regions that minimize formation of undesired byproducts (*25*). Consistent with the parental antibodies binding to distinct epitopes, both arms of CoV-X4042 can simultaneously engage the same SD1-RBD molecule (Fig. 5, A and B and fig. S8). In pseudovirus-based neutralization assays, CoV-X4042, like the sd1.040/rbd.042 cocktail, exhibited significant synergistic activity over either of the parental antibodies alone (Fig. 5C). CoV-X4042 neutralized pseudoviruses corresponding to VOC, and remained effective even when one of the parental antibodies lost efficacy (e.g. rbd.042 against Omicron BA.4/BA.5; Fig. 5D). Efficacy of the bispecific antibody was confirmed against ancestral and Omicron SARS-CoV-2 *in vitro* and by *in vivo* protection experiments showing that mice treated with CoV-X4042, either as pre- or post-exposure prophylaxis, maintained body weight and displayed diminished pathology and infectious virus titers in the lungs (Fig. 5, E to G). Therefore, bispecific antibodies composed of moieties that simultaneously target conserved neutralizing epitopes on SD1 and RBD are effective against SARS-CoV-2 in preclinical models.

## Discussion

The evolving antigenic landscape of SARS-CoV-2 poses unanticipated challenges to the development of vaccines and immunotherapies. Monoclonal antibodies against the RBD that are potent against the ancestral virus were prioritized for clinical development (*26–30*). However, most of these antibodies lost efficacy due to aa changes in SARS-CoV-2 variants that possibly resulted from immune pressure (*8, 10, 27, 31–33*). While antibodies have been reported to broadly-neutralize SARS-CoV-2 variants (e.g., anti-NTD antibodies, Bebtelovimab, Sotrovimab, *etcetera*), the majority of potent neutralizers recognize epitopes outside of coldspot regions (*6, 34*). Interestingly, the three RBD coldspots (residues 377-404, 418-438, and 454-476) partially comprise the conserved class 4 epitope, which has been identified as a potential target for eliciting cross-reactive antibodies against the sarbecovirus lineage (*35–39*). However, class 4 antibodies are weakly-neutralizing and lack effectiveness against Omicron sublineages (*40*). Neutralizing antibodies that target the S2 region and act in the post-attachment phase were recently described. Although generally weak neutralizers, some display broad reactivity against SARS-CoV-2 variants, and even against more distant CoV *species* (*16–20, 41, 42*), which is analogous to what is observed with antibodies that target the FP of influenza and HIV-1 (*43–45*).

In contrast, we discovered an antibody to the stem helix that precedes the HR2 region (hr2.016), which is at the same time potent (IC_50_ of 10 ng/mL) and broadly cross-reactive with beta- and also with some alpha- and gammacoronaviruses. Similarly, we identified a panel of anti-FP neutralizing antibodies that broadly recognize more distant coronaviruses, including all 9 of the known human CoVs. These results are consistent with the high degree of aa conservation at these regions of CoVs. It is noteworthy that IgG to FP and near the HR2 region were detected in convalescents and in individuals immunized with inactivated virus-based, but not with mRNA- or adenovirus-based vaccines, suggesting that not all the current vaccines are equally proficient at inducing antibodies to these broadly conserved neutralizing epitopes. This is possibly due to amino acid changes made in vaccines to stabilize the S trimer that may reduce the accessibility of these epitopes (*18*). Future vaccine development may consider including these new targets to afford improved and broader effectiveness. However, it will be important to understand how neutralizing antibodies recognize native epitopes on the prefusion spike trimer to guide such efforts.

Interestingly, for FP-specific antibodies, we observed specific contacts with residue R_815_, in the S2’ cleavage site. Cleavage at this site during entry allows for the rearrangement of the S2 subunit and subsequent fusion with the host cell membrane. A recent publication comparing CoV fusion peptide binding antibodies hypothesizes that binding to this residue may be a feature that distinguishes neutralizing and non-neutralizing fusion peptide antibodies (*19*). Another publication also posited that the mechanism of neutralization for cross-reactive FP antibodies was the steric occlusion of TMPRSS2 binding, rather than the prevention of fusogenic rearrangements, which supports the idea that binding to this highly-conserved Arg is a key determinant for neutralizing FP antibodies (*18*). In addition to TMPRSS2 cleavage, viral fusion can proceed through Cathepsin cleavage in the endosome downstream of the putative S2’ cleavage site (*2, 3*). Consistent with the observation that FP antibodies are effective regardless of TMPRSS2 expression on target cells, superposition of fp.006 on the S structure indicates that access to the Cathepsin site is also hindered by the antibody presence suggesting that fp.006 neutralization may involve a second mechanism preventing fusion of the viral and endosomal membranes in the cytoplasm.

Herein, we also report on human neutralizing antibodies that target SD1, a generally understudied but highly conserved region of S next to RBD. Previous studies have demonstrated that quaternary interactions between the RBD and SD1 of one protomer, with the NTD of the neighboring S1 protomer likely play a stabilizing role for prefusion S trimers (*46*). Our data suggest that antibody sd1.040 does not neutralize by interfering with ACE2 receptor binding. Instead, its function is likely linked to inhibition of conformational changes that expose the fusion loop upon engagement of ACE2. This is different from murine antibody S3H3 that potentially functions by “locking” the release of S1 subunits from S2 (*24*).

Even though the target sequences are conserved, FP, HR2 and SD1 antibodies are variably efficacious against VOC. This may reflect differences in the overall conformation of S variants that alter epitope accessibility, as well as changes in residues contacted by the antibodies that are outside of the coldspot region. For FP and HR2 antibodies, the observation that ACE2 attachment renders the epitopes more accessible, and improves neutralization, suggests that for therapeutic purposes it may be valuable to combine them with antibodies against the RBD that mimic ACE2 binding (*47*).

In recent work, broadly cross-reactive antibodies were discovered through large-scale screening of over 670,000 memory B cell antibodies or over 4000 B cell cultures (*18, 19*). By zooming in on portions of the virus that are both functionally relevant and averse to change, the strategy described herein represents a complementary, resource-savvy approach for the rapid discovery of antibodies with potential of being broadly cross-reactive against variants of a single virus or against multiple related virus *species*. This approach relies upon the analysis of large collections of virus sequences, which at present are publicly available only for a handful of pathogens (e.g. over 10 mio for SARS-CoV-2; 0.366 mio for influenza; 0.016 mio for HIV-1) (*14, 48, 49*), but are expected to become more broadly available for other pathogens in the future thanks to increased surveillance and ease of sequencing.

## Acknowledgments

We are grateful to all study participants and their families, as well as to the medical personnel of the Clinica Luganese Moncucco. We further thank Theodora Hatziioannou and Paul Bieniasz (Rockefeller University) for sharing plasmids and protocols for SARS-CoV-2 pseudovirus, Diego Morone, Irene Cassaniti, Josè Camilla Sammartino, Jan Kamis and Marketa Spevakova for technical assistance, Helena Jirincova (National Institute of Public Health, Prague) for providing virus isolates and Petra Strakova for help with virus cultivation, Nima Rezaei and Zahra Chavoshzadeh (Tehran University of Medical Sciences), for their contribution to the collection of vaccinated samples, and Pamela Bjorkman (California Institute of Technology) for sharing CoV expression plasmids. For the IMMUNO_COV study, we acknowledge the contribution of Francesca Montagnani, Massimiliano Fabbrini, Ilaria Rancan, Gabiria Pastore, Jacopo Polvere and Sara Zirpoli (University of Siena and University Hospital of Siena) for participants enrollment, clinical samples collection and processing. Cryo-EM data for this work was collected at the Stanford-SLAC cryo-EM center with support from Elizabeth Montabana. X-ray crystallographic data were collected at the Stanford Synchrotron Radiation Lightsource, SLAC National Accelerator Laboratory, which is supported by the U.S. Department of Energy, Office of Science, Office of Basic Energy Sciences under Contract No. DE-AC02-76SF00515. The SSRL Structural Molecular Biology Program is supported by the DOE Office of Biological and Environmental Research, and by the National Institutes of Health, National Institute of General Medical Sciences (P30GM133894). This study was supported in part by the Swiss Vaccine Research Institute (SVRI), George Mason University Fast Grant, Fondation Philanthropique Famille Sandoz, NIH grants U01 AI151698 (United World Antiviral Research Network, UWARN), P01 AI138938 and U19 AI111825 (to D.F.R.); European Union’s Horizon 2020 research and innovation programme under grant agreement no. 101003650 (to Q.P.-H, H.M, F.B., L.C., L.V. and D.F.R.); BRIDGE 40B2-0_203488 (to A.Ca. and D.F.R.); Department of Medical Biotechnologies, University of Siena (to D.M.); EU Joint Research Centre Exploratory Research program (project ‘FUTURE’, to G.G.); Swedish Research Council and Knut and Alice Wallenberg Foundation (to Q.P.-H.); Czech Ministry of Agriculture (RVO0518, to D.R.); National Institute of virology and bacteriology, Programme EXCELES, Project No. LX22NPO5103 funded by the European Union - Next Generation EU (to D.R.); Strategy of the Czech Academy of Sciences, AV21 - Virology and Antiviral Therapy (to D.R. and R.S.); Czech Science Foundation and Swiss National Science Foundation (21- 05445L; to D.R. and D.F.R.); 22-17139K by Czech Science Foundation (to R.S.); Czech Centre for Phenogenomics provided by Ministry of Education, Youth and Sports of the Czech Republic (LM2018126, to R.S.). A.Ca. thanks Mrs. Flora Gruner for her generous support. C.O.B. is supported by the Howard Hughes Medical Institute Hanna Gray Fellowship and is a Chan Zuckerberg Biohub investigator.

## Funding

Swiss Vaccine Research Institute (D.F.R.)

George Mason University Fast Grant (D.F.R.)

Fondation Philanthropique Famille Sandoz (D.F.R.)

National Institutes of Health grant U01AI151698, United World Antiviral Research Network - UWARN (D.F.R.)

National Institutes of Health grant P01AI138938 (to D.F.R.)

National Institutes of Health grant U19AI111825 (to D.F.R.)

European Union’s Horizon 2020 research and innovation programme grant no. 101003650, Antibody Therapy Against Coronavirus - ATAC (to Q.P.-H, H.M, F.B., L.C., L.V. and D.F.R.)

Swiss National Science Foundation grant BRIDGE 40B2-0_203488 (to A.Ca. and D.F.R.)

Department of Medical Biotechnologies, University of Siena (to D.M.)

EU Joint Research Centre Exploratory Research program project ‘FUTURE’ (to G.G.)

Swedish Research Council (to Q.P.-H.)

Knut and Alice Wallenberg Foundation (to Q.P.-H.)

Czech Ministry of Agriculture grant RVO0518 (to D.R.)

National Institute of virology and bacteriology, Programme EXCELES, Project No. LX22NPO5103 funded by the European Union - Next Generation EU (to D.R.)

Strategy of the Czech Academy of Sciences AV21, Virology and Antiviral Therapy (to D.R. and R.S.)

Czech Science Foundation and Swiss National Science Foundation grant 21-05445L (to D.R. and D.F.R.)

Czech Centre for Phenogenomics provided by Ministry of Education, Youth and Sports of the Czech Republic grant LM2018126 (to R.S.)

Howard Hughes Medical Institute Hanna Gray Fellowship (to C.O.B.)

Chan Zuckerberg Biohub (to C.O.B.)

U.S. Department of Energy, Office of Science, Office of Basic Energy Sciences under Contract No. DE-AC02-76SF00515

U.S. Department of Energy, Office of Biological and Environmental Research, and National Institutes of Health, National Institute of General Medical Sciences grant P30GM133894

## Author contributions

Conceived the study: F.Bi., V.Cr., A.Ca., C.O.B., D.F.R.

Designed and analyzed the experiments: F.Bi., V.Cr., M.E.A., C.Gu., M.Pa., M.Pe., L.C., R.S., D.R., L.V., A.Ca., C.O.B., D.F.R.

Designed human subjects research protocols: A.F.-P., M.U., D.F.R., C.Ga.

Produced reagents and carried out experiments: F.Bi., V.Cr., M.E.A., C.Gu., M.Pa., M.Pe., R.D.G., L.S., M.M., T.C.R., J.M., L.P., A.S., A.Ce., J.C., M.S., V.H., V.P., T.M., J.P., G.G., A.Ci., D.J., J.S., D.F., D. Meh., A.Ci. P.S., T.F.

Contributed reagents and samples: H.M., A.P., H.A., D.Med., Q.P.-H., F.Ba.

Recruited participants and executed human subjects research protocols: M.B., V.Ca., A.F.-P. Processed human samples: C.T.

Performed structural analysis: M.E.A., C.O.B.

Performed bioinformatic analysis: V.Cr., L.S., M.M, L.V., A.Ca.

Wrote the manuscript with input from all co-authors: F.Bi., V.Cr., M.E.A., C.O.B., D.F.R.

### Data and materials availability

Data supporting the findings of this study are available within the paper and its supplementary information files. All other data and materials are available from the corresponding authors upon reasonable request and may require MTA. Published data used in this study were taken from GenBank (https://www.ncbi.nlm.nih.gov/genbank/), UniProt (https://www.uniprot.org/), Protein Data Bank, PDB (https://www.rcsb.org/), Viralzone (https://viralzone.expasy.org/9556) and GISAID database (https://www.gisaid.org/). The script for coldspot identification is available at GitHub (https://github.com/cavallilab/coldspot). The atomic models and corresponding maps for all structures are available online under the following accession codes. For fp.006-FP, the model and structure factor files are available in the Protein Data Bank (PDB) under accession code 8D47. For sd1.040-S, the model and cryo-EM maps are available both in the PDB and Electron Microscopy Data Bank (EMDB) under codes 8D48 and EMD-27177, respectively.

## Supplementary Materials

Materials and Methods Figs. S1 to S8

Tables S1 to S4

## Materials and Methods

### Study participants

<u>COVID-19 convalescent cohort:</u> 67 individuals, who were diagnosed with COVID-19 at the Clinica Luganese Moncucco (CLM, Switzerland) between March and November of 2020, were enrolled in the study and written informed consent was obtained. Inclusion criteria were a SARS-CoV-2 positive nasopharyngeal swab test by real-time reverse transcription-polymerase chain reaction (RT-PCR) or being a symptomatic close contact (same household) of a hospitalized participant, and age ≥18 years. Samples were 83-269 days after onset of symptoms. The study was performed in compliance with all relevant ethical regulations under study protocols approved by the Ethical Committee of the Canton Ticino (ECCT): CE-3428 and CE-3960.

Control cohort: 17 individuals (≥18 years) with absence of prior SARS-CoV-2 infection or vaccination, as confirmed by negative serologic test, were enrolled between November 2020 and June 2021 and written informed consent was obtained (ECCT: CE-3428).

<u>Vaccination cohort:</u> individuals (≥18 years) with absence of prior SARS-CoV-2 infection and who received either mRNA-based (n=11 for BNT162b2, samples obtained 75-136 days after second dose; n=5 for mRNA-1273, 85-120 days after second dose), adenovirus-based (n=19 for ChAdOx1-S, 90 days after second dose; n=4 for Ad26.COV2.S, 21 days after single dose), or inactivated virus-based (n=2 for Sinovac, 26-60 days after second dose; n=24 for Sinopharm, 6-60 days after second dose) COVID-19 vaccines were enrolled under approved protocols (ECCT: CE-3428 and CE-3960; Ethic Committee Karolinska Institutet: Dnr 2020-02646; Ethic Committee Tehran University of Medical Sciences: IR.TUMS.CHMC.REC.1399.098-B2; Ethical Committee for Clinical Experimentation of Regione Toscana Area Vasta Sud Est [CEASVE]: ID 18869). Controls to the adenovirus vaccinated group are pre-vaccination samples from the same participants.

Pre-pandemic common cold coronavirus convalescents: 6 samples from individuals with confirmed common cold CoV infection were obtained 6-375 days after symptoms onset at Policlinico San Matteo, Pavia (Institutional Review Board protocol number P_20200029440).

#### Blood sample processing and storage

Peripheral blood mononuclear cells (PBMCs) from COVID-19 convalescents were obtained by Histopaque density centrifugation and stored in liquid nitrogen in the presence of FBS and DMSO. Anticoagulated plasma was aliquoted and stored at -20°C or less. Prior to experiments, aliquots of plasma were heat-inactivated (56°C for 1 hour) and then stored at 4°C. Similarly, frozen plasma aliquots from non-infected, common cold-infected, and vaccinated individuals were stored at 4°C after heat-inactivation.

### Peptides and recombinant proteins for biochemical studies

<u>Peptides:</u> Synthetic peptides containing the FP and HR2 coldspot sequences were designed and obtained (> 75% purity) from GenScript (Hong Kong). Peptides were biotinylated (biotin-Ahx) at the N-terminus and amidated at the C-terminus. The aa sequence of all peptides in this study is shown in fig. S2B.

Proteins: The CoV proteins were produced and purified as described (*50*).

S proteins: Codon-optimized gene encoding residues 1–1208 of SARS-CoV-2 S ectodomain (GenBank: MN908947) was synthesized and cloned into the mammalian expression vector pcDNA3.1(+) by Genscript; the sequence contains proline substitutions at residues 986 and 987 (S-2P), a ‘GSAS’ substitution at the furin cleavage site (residues 682–685), a C-terminal T4 fibritin trimerization motif and a C-terminal octa-histidine tag. SARS-CoV-2 S ectodomains corresponding to the SARS-CoV-2 VOC were based on: Delta, GenBank: QWK65230.1; Omicron BA.1 GenBank: UFO69279.1; Omicron BA.4/BA.5 GenBank: UPP14409.1 + G3V. MERS and HCoV-229E S ectodomains were based on PDB: 6NB3_A for MERS and PDB: 6U7H_A for HcoV-229E (residues 17-1142). RBD and SD1-RBD proteins: Plasmids for the production of RBD and SD1-RBD proteins were similarly designed and obtained. RBD and SD1-RBD corresponding to ancestral SARS-CoV-2 were based on an early SARS-CoV-2 sequence isolate (GenBank: QHO60594.1), and included aa 331-529 and 319-591, respectively. RBD and SD1-RBD corresponding to the VOC were based on: Alpha, GenBank: QWE88920.1; Beta, GenBank: QRN78347.1; Gamma, GenBank: QVE55289.1; Delta, GenBank: QWK65230.1; Omicron BA.1, GenBank: UFO69279.1; Omicron BA.2, GenBank: UJE45220.1; Omicron BA.2.75, GenBank: UTM82166.1; Omicron BA.2.75.2, GenBank: UTM82166.1 + R343T + F483S; Omicron BA.4/BA.5, GenBank: UPP14409.1; Omicron BA.4.6, GenBank: UPP14409.1 + R341T; Omicron BA.4.7, GenBank: UPP14409.1 + R341S; Omicron BA.5.9, GenBank: UPP14409.1 + R341I. SD1-RBD T572I and E583D were based on the ancestral SARS-CoV-2 sequence (GenBank: QHO60594.1) with T572I or E583D, respectively. For flow cytometry-based sorting experiments, ancestral SARS-CoV-2 RBD and SD1-RBD constructs were produced that included at the C-terminus an Avi-tag (GLNDIFEAQKIEWHE) for site-directed biotinylation in addition to octa-histidine tag for purification. ACE2 protein (human ACE2 fused at the C-terminus with the Fc of mouse IgG) was as previously (*50*), with synthetic, codon-optimized nucleotide sequence of hACE2 (residues 18- 740) fused at the C-terminus to the Fc region of human IgG1 and cloned into pcDNA3.1(+) vector by Genscript.

All proteins were produced by transient PEI transfection in Expi293F cells (ThermoFisher), purified from the cell supernatants with proper affinity columns and analyzed to ensure functionality, stability, lack of aggregation and batch-to-batch reproducibility as previously described (*50*).

#### ELISAs

To evaluate the ability of antibodies to bind to peptides and proteins of CoVs, we performed enzyme-linked immunosorbent assays (ELISA).

<u>Peptide ELISA</u>: 96-well plates (ThermoFisher, 442404) were coated with 50 μl per well of a 2μg/ml Neutravidin (Life Technologies, 31000) solution in PBS, overnight at room temperature. Plates were washed 4 times with washing buffer (PBS + 0.05% Tween-20 [Sigma-Aldrich]) and incubated with 50 μl per well of a 50 nM biotinylated peptide solution in PBS for 1 h at room temperature. After washing 4 times with washing buffer, plates were incubated with 200 μl per well of blocking buffer (PBS + 2% BSA + 0.05% Tween-20) for 2 h at room temperature. Plates were then washed 4 times with washing buffer, and serial dilutions of monoclonal antibodies or plasma were added in PBS + 0.05% Tween-20 and incubated for 1 h at room temperature. To screen for the presence of anti-coldspot peptide IgGs, plasma samples were assayed at 1:50 starting dilution followed by 7 (Fig. 1C and 3B) or 3 (fig. S1B) threefold serial dilutions. Monoclonal antibodies were tested starting at the indicated concentrations and followed by threefold serial dilutions. Plates were subsequently washed 4 times with washing buffer and incubated with anti-human IgG secondary antibody conjugated to horseradish peroxidase (HRP) (GE Healthcare, NA933) at a 1:5000 dilution in PBS + 0.05% Tween-20. Finally, after washing 4 times with washing buffer, plates were developed by the addition of 50 μl per well of the HRP substrate TMB (ThermoFisher, 34021) for 10 min. The developing reaction was stopped with 50 μl per well of a 1M H_2_SO_4_ solution, and absorbance was measured at 450 nm with an ELISA microplate reader (BioTek) with Gen5 software. The Area Under the Curve (AUC) was obtained from two independent experiments and plotted with GraphPad Prism.

<u>Protein ELISA</u>: Experiments were performed with 96-well plates coated with 50 μl per well of a 5 μg/ml protein solution in PBS overnight at room temperature, and subsequently blocked and treated as described above. Monoclonal antibodies were tested starting at the indicated concentrations and followed by three-, four- or fivefold serial dilutions. Cross-reactivity ELISAs on SD1-RBD variants were performed in 96-well plates with half-area (Corning, 3690) and using half of the volumes mentioned above.

<u>ACE2 binding ELISA:</u> Experiments were as previously described (*50*). Briefly, 96-well plates with half-area (Corning, 3690) were coated with 25 μl per well of a 5 μg/mL spike solution in PBS and incubated overnight at 4°C. After washing, blocking was performed with 10% FBS in PBS for 1 h at RT. Monoclonal antibodies were added at the indicated concentrations and followed by threefold serial dilutions in blocking buffer. After washing, 25 μl per well of a 5 μg/mL solution of human ACE2 fused to the Fc portion of mouse IgG were added to the plate. Detection of ACE2 was performed with an AP-conjugated anti-mouse IgG secondary Ab (Southern Biotechnology Associates, 1030-04) diluted 1:500 in blocking buffer.

#### Protein biotinylation for use in flow cytometry

Purified, Avi-tagged SARS-CoV-2 RBD and SD1-RBD (both corresponding to SARS-CoV-2 ancestral virus) were biotinylated using the Biotin-Protein Ligase-BIRA kit according to manufacturer’s instructions (Avidity). Ovalbumin (Sigma, A5503-1G) was biotinylated using the EZ-Link Sulfo-NHS-LC-Biotinylation kit according to the manufacturer’s instructions (Thermo Scientific). Biotinylated Ovalbumin and RBD were conjugated to streptavidin-BV711 (BD biosciences, 563262) and SD1-RBD to streptavidin-PE (BD biosciences, 554061) and streptavidin-Alexa Fluor 647 (AF647, Biolegend, 405237), respectively.

#### Single-cell sorting by flow cytometry

B cells from PBMCs of uninfected controls or of COVID-19 convalescent individuals were enriched using the pan-B-cell isolation kit according to manufacturer’s instructions (Miltenyi Biotec, 130-101-638). The enriched B cells were subsequently stained in FACS buffer (PBS + 2% FCS + 1mM EDTA) with the following antibodies/reagents (all at 1:200 dilution) for 30 min on ice: anti-CD20-PE-Cy7 (BD Biosciences, 335828), anti-CD14-APC-eFluor 780 (Thermo Fischer Scientific, 47-0149-42), anti-CD16-APC-eFluor 780 (Thermo Fischer Scientific, 47-0168-41), anti-CD3-APC-eFluor 780 (Thermo Fischer Scientific, 47-0037-41), anti-CD8-APC-eFluor 780 (Invitrogen, 47-0086-42), Zombie NIR (BioLegend, 423105), as well as fluorophore-labeled ovalbumin (Ova) and peptides. Live single Zombie-NIR^−^CD14^−^CD16^−^CD3^−^CD8^−^CD20^+^Ova^−^peptide-PE^+^peptide-AF647^+^ B cells were single-cell sorted into 96-well plates containing 4 μl of lysis buffer (0.5× PBS, 10 mM DTT, 3,000 units/ml RNasin Ribonuclease Inhibitors [Promega, N2615]) per well using a FACS Aria III, and the analysis was performed with FlowJo software. The isolation of SD1-enriched B cells was performed similarly, except that sorted cells were live single Zombie-NIR^−^CD14^−^CD16^−^CD3^−^CD8^−^CD20^+^Ova^−^RBD^−^SD1-RBD-PE^+^SD1-RBD-AF647^+^. The gating strategy is shown in fig. S1C.

#### Antibody gene sequencing, cloning and expression

Antibody gene sequences were identified as described previously (*4*). Briefly, single cell RNA was reverse-transcribed (SuperScript III Reverse Transcriptase, Invitrogen, 18080-044) and the cDNA stored at -20°C or used for subsequent amplification of the variable IGH, IGL and IGK genes by nested PCR and Sanger sequencing. Amplicons from the first PCR reaction were used as templates for Sequence- and Ligation-Independent Cloning (SLIC) into antibody expression vectors. Recombinant monoclonal antibodies and Fabs were produced and purified as previously described (*51*). C121, C135 anti-SARS-CoV-2 antibodies and isotype control anti-Zika virus antibody Z021 were previously published (*4, 51*); the sequences of antibodies CV3-25 and S2P6 were derived from the literature (GenBank: MW681575.1 and MW681603.1 (*17*); PDB 7RNJ (*16*)) and produced in house starting from synthetic DNA (Genscript). The human IgG-like bispecific CoV-X4042 was designed based on the variable regions of antibodies sd1.040 and rbd.042 in the CrossMAb format (*25*). Four pcDNA3.1(+) mammalian expression plasmids for CrossMAb production were synthesized (Genscript), used to transfect Expi293F cells (ThermoFisher) in a 1:1:1:1 ratio, and purified from the cell supernatants as previously described (*50*). All the antibodies were tested to ensure functionality, stability and batch-to-batch reproducibility.

#### SARS-CoV-2 pseudotyped reporter viruses

The generation of plasmids to express a C-terminally truncated SARS-CoV-2 S protein (pSARS-CoV2-S_trunc_), the HIV-1 structural/regulatory proteins (pHIV_NL_GagPol) and the NanoLuc reporter construct (pCCNanoLuc2AEGFP) were previously described (*52*), and like the pSARS-CoV2-S_trunc_ plasmid for Delta variant, they were kindly gifted by Drs. Paul Bieniasz and Theodora Hatziioannou (The Rockefeller University, New York). Plasmids expressing Alpha, Beta, Gamma, Delta, Omicron BA.1, BA.2, BA.2.75, BA.2.75.2 and BA.4/BA.5. SARS-COV-2-S_trunc_ variants were generated in house by site-directed mutagenesis (QuikChange Multi Site-Directed Mutagenesis Kit, Agilent) starting from synthetic DNA (Genscript). The sequences corresponding to SARS-CoV-2 VOC were based on: Alpha (B1.1.7; GenBank QWE88920.1), Beta (B.1.351; GenBank QRN78347.1), Gamma (B.1.1.28.1; GenBank QRX39425.1), Delta (B.1.617.2; GenBank QWK65230), Omicron BA.1 (B.1.1.529; GenBank UFO69279.1), Omicron BA.2 (GenBank ULB15050.1), Omicron BA.2.75 (GenBank UTM82166.1), Omicron BA.2.75.2 (GenBank UTM82166.1 + R343T + F483S + D1196N), Omicron BA.4/BA.5 (GenBank UPP14409.1 + G3V). In all pseudoviruses, the intracellular domain was similarly truncated and the S1/S2 furin cleavage site was unchanged. The generation of pseudotyped virus stocks was as previously described, with minor modifications (*51*). Briefly, 293T cells were transfected with pHIV_NL_GagPol, pCCNanoLuc2AEGFP and pSARS-CoV2-S_trunc_ plasmids using PEI-MAX (Polysciences). At 24h after transfection, supernatants containing non-replicating virions were harvested, filtered and stored at -80°C. Infectivity was determined by titration on 293T_ACE2_ and 293T_ACE2/TMPRSS2_ cells.

#### Pseudotyped virus neutralization assay

The assay was previously described (*52*). Briefly, three- or four-fold serially diluted monoclonal antibodies were incubated with the SARS-CoV-2 pseudotyped virus for 1 hour at 37°C degrees. The mixture was subsequently incubated with 293T_ACE2_ or 293T_ACE2/TMPRSS2_ cells for 48 hours, after which cells were washed once with PBS and lysed with Luciferase Cell Culture Lysis 5x reagent (Promega). Nanoluc Luciferase activity of lysates was then measured using the Nano-Glo Luciferase Assay System (Promega) with GloMax Discover System reader (Promega). Relative luminescence units were then normalized to those derived from cells infected with SARS-CoV-2 pseudotyped virus in the absence of monoclonal antibodies. The half-maximal inhibitory concentration of monoclonal antibodies (IC_50_) was determined using four-parameter nonlinear regression curve fit (GraphPad Prism).

#### Detection of monoclonal antibody binding to S by flow cytometry

1.9x10_6_ 293T cells were plated in 60x15 mm dishes (Corning, Ref#430166) and co-transfected with two plasmids encoding GFP (2.25μg) and the SARS-CoV-2 S protein (pSARS-CoV2-S_trunc_; 2.25μg) using 18 μg PEI-MAX as a transfection reagent 24 hours later. 40 hours upon transfection, cells were collected by gentle pipetting, and 50’000 transfected cells per well (in U-bottom 96-well plates; Corning, Ref#3799) were subsequently stained with 10 μg/ml pre-labelled monoclonal antibodies in the presence or absence of 30 μg/ml human ACE2 in a total volume of 100 μl PBS supplemented with 5% FBS and 2 mM EDTA for 2h at room temperature similar to a previous report (*18*) (Fig. 2H). In Fig. 4F, cells were pre-incubated with sd1.040 or Z021 (isotype) antibodies at final concentration of 10 μg/ml for 30 minutes at room temperature before the addition of ACE2 and pre-labelled fp.006. Fluorescent labeling of monoclonal antibodies was performed with the DY-647P1-NHS-ester reagent (Dyomics, Ref#647P1-01) according to manufacturer’s instructions. After staining, cells were washed twice, acquired with BD FACSCanto and analyzed with FlowJo software.

#### Inhibition of cell-cell fusion

Inhibition of spike-mediated cell-cell fusion was tested using an assay developed by Invivogen. Briefly, hMyD88 expressing 293 cells (Invivogen, cat. code 293-hmyd) were transfected with Wuhan pSARS-CoV2-S_trunc_ plasmid using jetOptimus® (Polyplus). At 24h after transfection, cells were resuspended in complete media and incubated with 200μg/mL of antibodies for 1h at 37°C, before addition of SEAP reporter 293 cells expressing hACE2 (Invivogen, cat. code hkb-hace2). Cells were co-cultured for 24h and cell-cell fusion was assessed measuring secreted embryonic alkaline phosphatase (SEAP) activity into cells supernatant using QUANTI-Blue™ Solution (Invivogen), according to the manufacturer’s protocol.

#### Surface Plasmon Resonance (SPR) assays

The IgG antibody or Fab binding properties were analyzed at 25 °C using a Biacore 8K instrument (GE Healthcare) with 10 mM HEPES pH 7.4, 150 mM NaCl, 3 mM EDTA and 0.005% Tween-20 as running buffer. SARS-CoV-2 antigens (SD1-RBD or S-2P) were immobilized on the surface of CM5 chips (Cytiva) through standard amine coupling. Increasing concentrations of IgG/Fab were injected using a single-cycle kinetics setting and dissociation was followed for 10 minutes. Analyte responses were corrected for unspecific binding and buffer responses. Curve fitting and data analysis were performed with Biacore Insight Evaluation Software v.2.0.15.12933. Competition experiments were performed to obtain information on the IgG/Fab binding regions. First, antibody was immobilized on the surface of CM5 chips (Cytiva) through standard amine coupling; SD1-RBD was then flowed to form SD1-RBD/antibody complex and, shortly thereafter, the second antibody was injected. If a binding event is detected at the final step, then the 2^nd^ antibody has a different epitope compared to the 1st (immobilized) antibody. If no binding event is detected, the two antibodies share overlapping epitopes. Competition experiments were also used to confirm the functionality of both arms of the CoV-X4042 bispecific. First, sd1.040 or rbd.042 antibodies were immobilized on the surface of CM5 chips (Cytiva) through standard amine coupling; then SD1-RBD was flowed to form RBD-SD1/antibody complex and, shortly thereafter, CoV-X4042 was injected. The analysis and comparison of kinetics parameters at different SD1-RBD concentrations were also performed as previously described (*49*) to assess the ability of CoV-X4042 to bind bivalently to a single SD1-RBD molecule.

#### Cell lines

293T_ACE2/TMPRSS2_ cell line was generated by transfecting 293T_ACE2_ (*51*) cells with pCMV3-FLAG-TMPRSS2 (SinoBiological) using Lipofectamine 3000 (Invitrogen) and selected with 200μg/mL Hygromycin B (Invivogen) two days post-transfection. 293T cells for pseudotyped virus production were cultured in DMEM supplemented with 10% FBS. 293T_ACE2_ cells were cultured in DMEM supplemented with 10% FBS, 1% NEAA, 1mM Sodium Pyruvate, 1x Penicillin/Streptomycin and 5μg/mL Blasticidin. Vero cells were from ATCC (CCL-81), Expi293F and 293FT from ThermoFisher (#A14528 and R70007). hMyD88 expressing 293 cells (Invivogen) were cultured in DMEM supplemented with 10% FBS, 1x Penicillin/Streptomycin and 10μg/mL Puromycin. SEAP reporter 293 cells expressing hACE2 (Invivogen) were grown in DMEM supplemented with 10% FBS, 1x Penicillin/Streptomycin, 1μg/mL Puromycin and 100μg/mL Zeocin.

#### Focus Reduction Neutralization Tests (FRNT)

The assay was performed similar to how previously described(*4*). Briefly, the day before infection, Vero cells were seeded at 1x10^4^ cells/well in 96-well plates. The antibodies were diluted to final concentrations in Dulbecco’s modified Eagle’s medium (DMEM) supplemented with 10% newborn calf serum, 100 U/mL penicillin, 100 µg/mL streptomycin, and 1% glutamine (Sigma-Aldrich, Prague, Czech Republic). Subsequently, the diluted samples were mixed with 1000 PFU/well of ancestral SARS-CoV-2 (strain SARS-CoV-2/human/Czech Republic/951/2020) or Omicron (B.1.1.529-like; hCoV-19/Czech Republic/KNL_2021-110119140/2021) and incubated at 37°C for 90 minutes. The antibody-virus mixture was then applied directly to Vero cells (MOI of ∼0.1 PFU/cell) and incubated for 22 hours at 37°C and 5% CO_2_. Cells were then fixed by cold acetone-methanol fixation (1:1, v/v) and blocked with 10% fetal bovine serum. Cells were incubated with a rabbit (2019-nCoV) S1 antibody (1:50; Sino Biological, Duesseldorf, Germany) and then incubated for 1 hour at 37°C with secondary goat anti-rabbit antibodies conjugated with fluorescein isothiocyanate (FITC; 1:250; Sigma-Aldrich, Prague, Czech Republic). Fluorescence intensity was measured using the Synergy H1 microplate reader (BioTek) with the following parameters: Plate type (96 WELL PLATE), fluorescence (area scan) excitation 490/emission 525, optics (Top) and gain (125), light source (Xenon Flash), lamp energy (High), reading speed (Normal), delay (100 msec), and reading height (6 mm). For Omicron, fluorescent foci were manually counted using an Olympus IX71 epifluorescence microscope and the numbers obtained normalized to no antibody control.

#### *In vivo* protection experiments

This study was performed in strict accordance with Czech laws and guidelines on the use of experimental animals and the protection of animals against cruelty (Animal Welfare Act No. 246/1992 Coll.). The protocol was approved by the Ethics Committee for Animal Experiments of the Institute of Parasitology, Institute of Molecular Genetics of the Czech Academy of Sciences, and by the Departmental Expert Committee for Approval of Projects of Experiments on Animals of the Czech Academy of Sciences (approvals 82/2020 and 101/2020). Thirteen-to fifteen-week-old female C57BL/6NCrl mice were ACE2-humanized by inhalation of a modified adeno-associated virus (AAV) (AAV-hACE2), as described previously (*50*). At least 7 days after application of AAV-hACE2 virus particles, mice were intranasally infected with SARS-CoV-2 (1 × 10^4^ plaque-forming units; ancestral strain SARS-CoV-2/human/Czech Republic/951/2020, or Omicron B.1.1.529-like; hCoV-19/Czech Republic/KNL_2021-110119140/2021, both isolated from clinical specimens at the National Institute of Public Health, Prague; passaged five times (six times for Omicron) in Vero E6 cells before use in this study) in a total volume of 50 μl DMEM. Twenty-four hours before (pre-exposure prophylaxis), or 2 hours after (post-exposure prophylaxis) infection, mice were injected intraperitoneally with either hr2.016, CoV-X4042 (both at 300 μg), fp.006 (500 μg in pre-exposure or 300 ug in post-exposure) or isotype control (either at 300 or 500 μg). Mice were culled at the indicated time points after infection, and their tissues were collected for analysis. Lung tissue was homogenized using Mixer Mill MM400 (Retsch, Haan, Germany) and processed as a 20% (w/v) suspension in DMEM containing 10% newborn calf serum. Homogenates were clarified by centrifugation at 14,000g (10 min, 4°C), and supernatant medium was used for plaque assay as previously described (*50*).

No sample-size calculation was performed. The sample sizes were chosen based on experience and previously published papers (e.g., (*53, 54*)). Details about groups and sample sizes for mouse virus challenge studies are provided in the figure legends. Experiments were successfully repeated at least twice. No data were excluded. The mice were randomly assigned to cages and the cages were then randomized into groups. Blinding was not relevant to this study. The readouts of all experiments could be assessed objectively. Mouse weight loss was determined using body weight measurement as a readout, and FRNT was used to quantify viral burden.

#### Computational analyses of viral sequences

Sequences of reference for ancestral SARS-CoV-2 and its variants of interest and concern (Fig. 1A) were derived from Viralzone (https://viralzone.expasy.org/9556), matched the WHO classification (https://www.who.int/en/activities/tracking-SARS-CoV-2-variants/) and were as follow: ancestral SARS-CoV-2 Wuhan-Hu-1 (19A; GenBank: QHO60594.1); Alpha (B1.1.7; GenBank: QWE88920.1); Beta (B.1.351; GenBank: QRN78347.1); Gamma (B.1.1.28.1; GenBank: QVE55289.1); Delta (B.1.617.2; GenBank: QWK65230.1); Epsilon (B.1.427; B.1.429; GenBank QQM19141.1); Zeta (B.1.1.28.2; GenBank; QQX30509.1); Eta (B.1.525; GenBank: QRF70806.1); Theta (B.1.1.28.3); Iota (B.1.526; GenBank: QRX49325.1); Kappa (B.1.617.1; GenBank: QTY83052.1); Lambda (B.1.1.1.C37; GenBank: QTJ90974.1); Mu (B.1.621); Omicron BA.1 (previously B.1.1.529; GenBank: UFO69279.1); Omicron BA.2 (GenBank: ULB15050.1); Omicron BA.3 (GISAID: EPI ISL 9092427); Omicron BA.4 (GenBank: UPP14409.1); omicron BA.5 (GenBank: UOZ45804.1). For the analysis of SARS-CoV-2 amino acid substitutions, viral sequences of S that were present in GISAID as of either 31 December 2020, 31 December 2021, or April 29 2022, were downloaded. Sequences with a length of S corresponding between 1223 and 1323 aa, and no undetermined aa, were aligned to determine the frequency of aa changes over ancestral reference sequence using BLASTP version 2.5.0 with default settings. Frequencies were computed using in house developed bash and C++ pipeline available at GitHub (https://github.com/cavallilab/coldspot). The models of the full S, glycosylated and with a membrane, both closed and open conformation, were taken from a previous publication (*55*) and rendered with Pymol 2.3.5 (Figs. 1B and 3A). For the phylogenetic analysis and peptide sequence alignment, sequences of representative S protein of CoV *species* classified according to the ICTV taxonomical classification (https://talk.ictvonline.org/taxonomy/) (*56*) were derived from the NCBI taxonomy database (https://www.ncbi.nlm.nih.gov/data-hub/taxonomy/11118/) (*57*), aligned using ClustalW (SnapGene), and the phylogenetic tree was built using phylogeny.fr with default settings (*58*).

#### X-ray crystallography

fp.006 Fab in 1X TBS (20 mM Tris, 150 mM NaCl) was mixed with the fusion peptide (PBS with 10% DMSO; PSKRSFIEDLLFNKVTLADA with N-terminal Biotin-Ahx and C-terminal amidation) at a 1:2 molar ratio (Fab:peptide). The sample was incubated overnight at room temperature, and then concentrated to ∼8.8 mg/mL using an Amicon spin filter with a 30 kDa molecular weight cutoff (Millipore Sigma) after diluting with an additional sample volume of 1X TBS to decrease the proportion of PBS and DMSO in the complex. Crystallization trials were set up using the sitting drop vapor diffusion method by mixing equal volumes of fp.006-FP complex and reservoir using a Douglas Oryx8 robot and commercially available 96-well crystallization screens (Hampton Research). Crystals were grown at 16 °C and observed in multiple conditions. The single crystal that was used for structure determination of fp.006-FP was obtained in 0.2 M Potassium phosphate monobasic pH 4.8 and 20% w/v Polyethylene glycol 3350, and were cryoprotected in a solution matching the reservoir and 30% glycerol and then cryocooled in liquid nitrogen.

X-ray diffraction data was collected at the Stanford Synchrotron Radiation Lightsource (SSRL) beamline 12-1 with an Eiger X 16 M pixel detector (Dectris) at a wavelength of 0.979 Å and temperature of 100 K. Data from a single crystal was indexed and integrated in XDS/Dials (*59*), and then merged using AIMLESS in CCP4 (*60*). Structures were determined using molecular replacement in PHASER (*61*) using two copies of each of the following individual chains as search models: V_H_ (PDB: 4GXU chain M with CRH3 trimmed), V_L_ (PDB: 6FG1 chain B with CDRL3 trimmed), C_H_ (PDB: 4GXU), and C_L_ (PDB: 6FG1). Coordinates were refined using iterative rounds of automated and interactive refinement in Phenix (*62*) and Coot (*63*), respectively. The final model contains 97.9% Ramachandran favored residues, with 2.0% allowed and the remaining 0.1% Ramachandran outliers.

#### Cryo-EM sample preparation

Concentrated and purified sd1.040 Fab was mixed with SARS-CoV-2 S-2P trimer at a 1.1:1 molar ratio (Fab:trimer) to a final concentration of 3 mg/mL and incubated at room temperature for thirty minutes. Immediately prior to application of 3.1 µL of sample to a freshly glow-discharged 300 mesh Quantifoil R1.2/1.3 grid, fluorinated octyl-maltoside (FOM) was added to a final concentration of 0.02% (w/v). Complex was vitrified by plunging into 100% liquid ethane after blotting for 3.5 s with Whatman No. 1 filter paper at 22 °C and 100% humidity using a Mark IV Vitrobot (Thermo Fisher).

#### Cryo-EM data collection and processing

Single particle movies were collected on a Titan Krios TEM (300 kV) using SerialEM automated data collection software (*64*) and a K3 camera (Gatan) behind a BioQuantum energy filter (Gatan) with a 20 eV slit size (0.85 Å/pixel). Specific collection parameters are summarized in table S4. Data processing followed a similar workflow as previously described (*65*). Briefly, 9,894 movies were patch motion corrected for beam-induced motion including dose-weighting within cryoSPARC v3.1 (*66*). CTF estimates were performed on non-doseweighted micrographs, which were subsequently curated to remove poor fits and images with thick ice. An initial set of particles were generated using automated blob picking, of which, a subset (300,000 particles) was used to generate four *ab initio* volumes. The entire particle stack (4,343,219 particles) were extracted 4x-binned and heterogeneously refined into the four *ab initio* volumes. Particles corresponding to the volume that best demonstrated features of a Fab bound to a protomer were cleaned up using 2D classification, and reextracted with 2x-binning. The resulting particle stack (1,552,774 particles) was further 3D classified in cryoSPARC (k=6). Particles corresponding to the 3D volumes with well-defined secondary structural features were pooled and homogeneously refined with C1 symmetry. Following CTF refinement and application of a mask for focus refinement on the RBD-SD1-Fab regions, the final reconstructed volume achieved a global resolution of 3.7Å based on gold standard FSC calculations.

#### Cryo-EM structure modeling and refinement

Coordinates for the sd1.040-S1 protomer complex were generated by docking individual domains from reference structures (individual spike domains, PDB 6VXX – chain A; Fab heavy chain, PDB 5AZE-chain H; Fab light chain, PDB 7D0D-chain L) into cryo-EM density using UCSF Chimera (*67*). Initial models were rigid body refined into cryo-EM density, followed by real space refinement with morphing in Phenix. Sequence matching was interactively performed in Coot and models were further refined in Phenix. Validation of model coordinates was performed using MolProbity (*68*).

#### Structural analyses

CDR and somatic mutation assignments were produced by IMGT V-QUEST (*69*). Graphics describing structures were made in ChimeraX (*70*). Buried surface areas were calculated using the online PDBePISA server (*71*). Contacting residues are defined as those with less than 4 Å distance between atoms of different chains. Hydrogen bond assignments were made using a 3.5 Å cutoff and A-D-H angle greater than 90°. RMSD calculations were done in PyMOL (Schrödinger). Antibody residues are numbered according to the Kabat convention.

#### Statistical analyses

Statistical significance between two groups of mice was determined using non-parametric two-tailed Mann–Whitney U-tests. For paired samples, we used two-tailed t-test. For P values calculation of Fig. 5C we used the more stringent Welch’s t-test, two-tailed, which does not assume equal variants of the 2 samples. Correlation between plasma IgG reactivity to SARS-CoV-2 RBD and SD1-RBD was assessed using Pearson correlation analysis. A p-value of less than 0.05 was considered statistically significant. In the figures, significance is shown as follow: ns p≥0.05 (not significant), *p<0.05, **p<0.01, ***p<0.001 and ****p<0.0001. Data and statistical analyses were performed with GraphPad Prism (version 8.4.3).

---

**Fig. S1.**
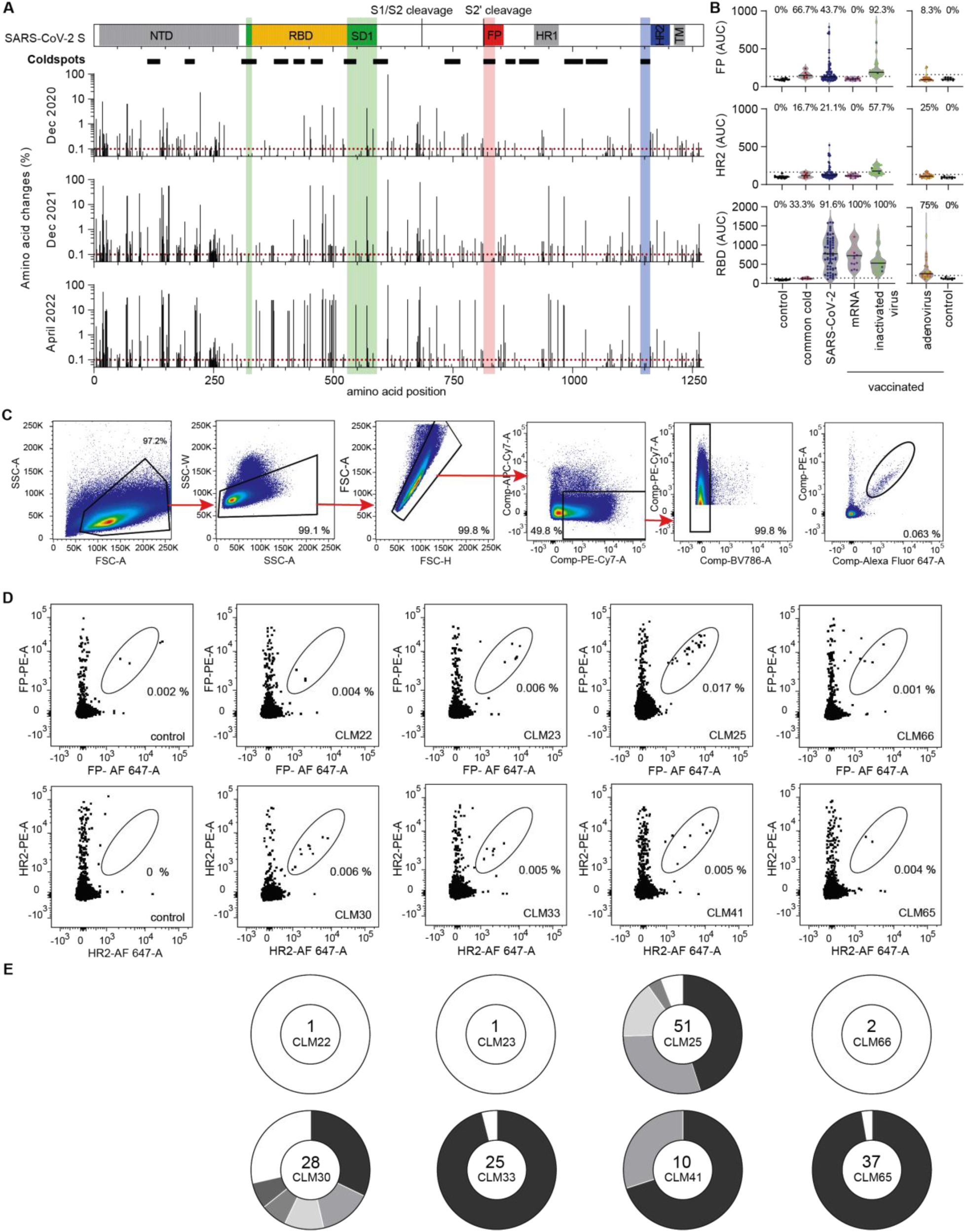
Discovery of coldspot antibodies. **(A)** Cartoon diagram of the SARS-CoV-2 S with highlighted coldspot regions at the fusion peptide (FP, red) and near heptad repeat 2 (HR2, blue), as well as subdomain 1 (SD1, green). The thick horizontal lines underneath indicate the location of all coldspots, and the histograms show the frequency of aa changes throughout S at different time points: 31.12.2020 (671, 006), 31.12.2021 (7,526,116) and 29.04.2022 (10,480,461 sequences from GISAID). **(B)** ELISA reactivity of plasma IgG antibodies to FP (top) and HR2 (middle) peptides or to SARS-CoV-2 RBD (bottom). SARS-CoV-2 (n=67) and common cold CoV (n=6) convalescent samples are compared to COVID-19 vaccinees (mRNA: BNT162b2 n=11, purple; mRNA-1273 n=5, pink; inactivated virus: Sinovac n=2, dark green; Sinopharm n=24, light green; adenovirus: ChAdOx1-S n=19, orange; Ad26.COV2.S n=4, yellow), and to non-infected controls (n=25). Area Under the Curve (AUC); average of two experiments. The percentage of samples with antibody levels 4 standard deviations above controls is shown on top. **(C)** Gating strategy for sorting peptide-specific or SD1-enriched B cells by flow cytometry. **(D)** Identification of FP-peptide specific (top) and HR2-peptide specific (bottom) B cells by flow cytometry. Numbers indicate percentage of gated double-positive cells. **(E)** Clonal analysis of antibody sequences derived from peptide-specific B cells identified in **(D)**. Pie charts show the total number of antibody sequences (center); the size of the slices is proportional to the number of clonally related sequences. White slices indicate antibody sequences that are not part of a clone.

**Fig. S2.**
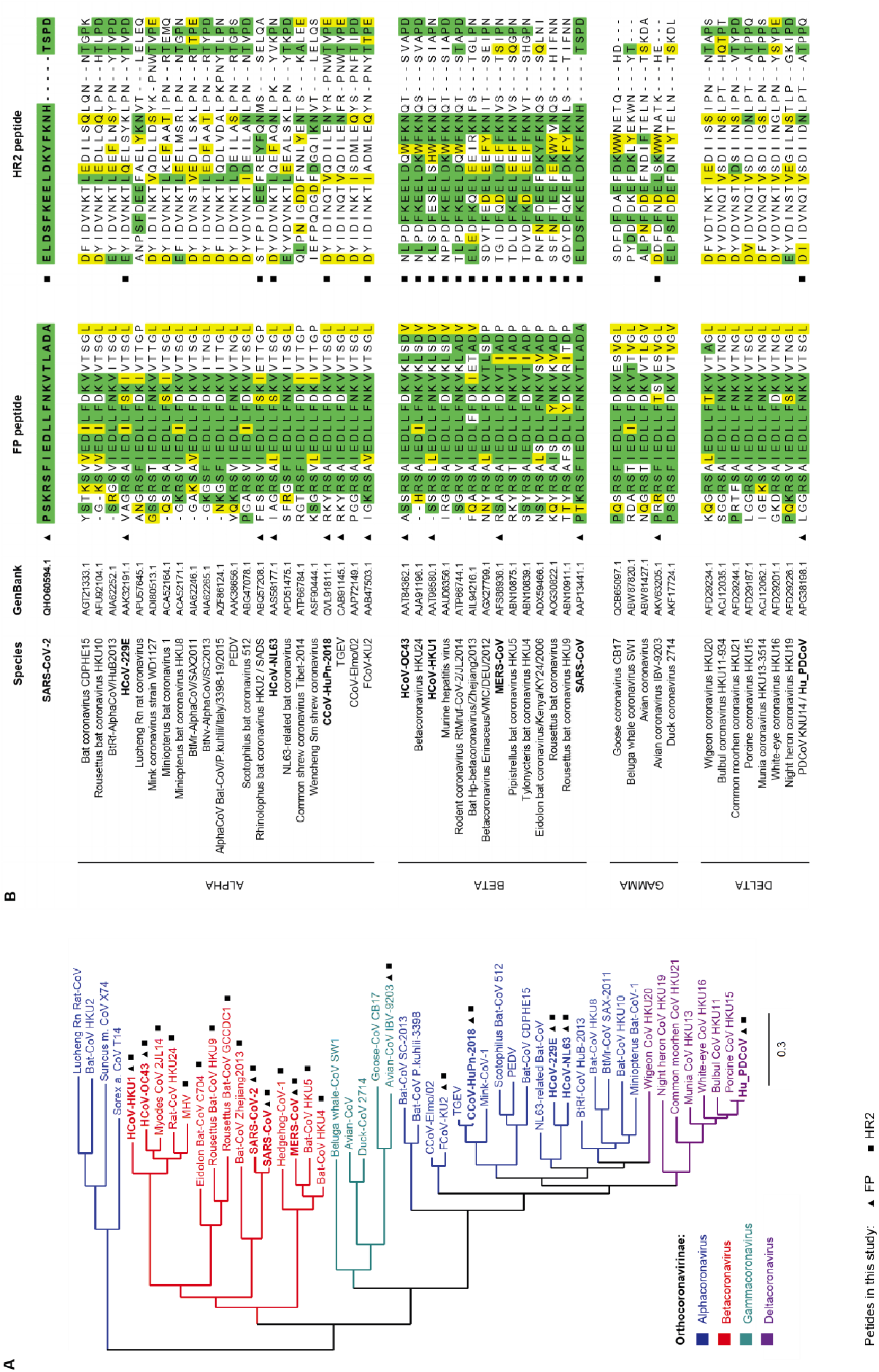
Similarity of peptides at the FP and HR2 of *Orthocoronavirinae*. **(A)** Phylogenetic analysis of CoV species based on the aa sequence of S. **(B)** Alignment of SARS-CoV-2 FP and HR2 peptide sequences with those corresponding to related CoVs. In green identical residues; in yellow residues that are similar by side chain functionality.

**Fig. S3.**
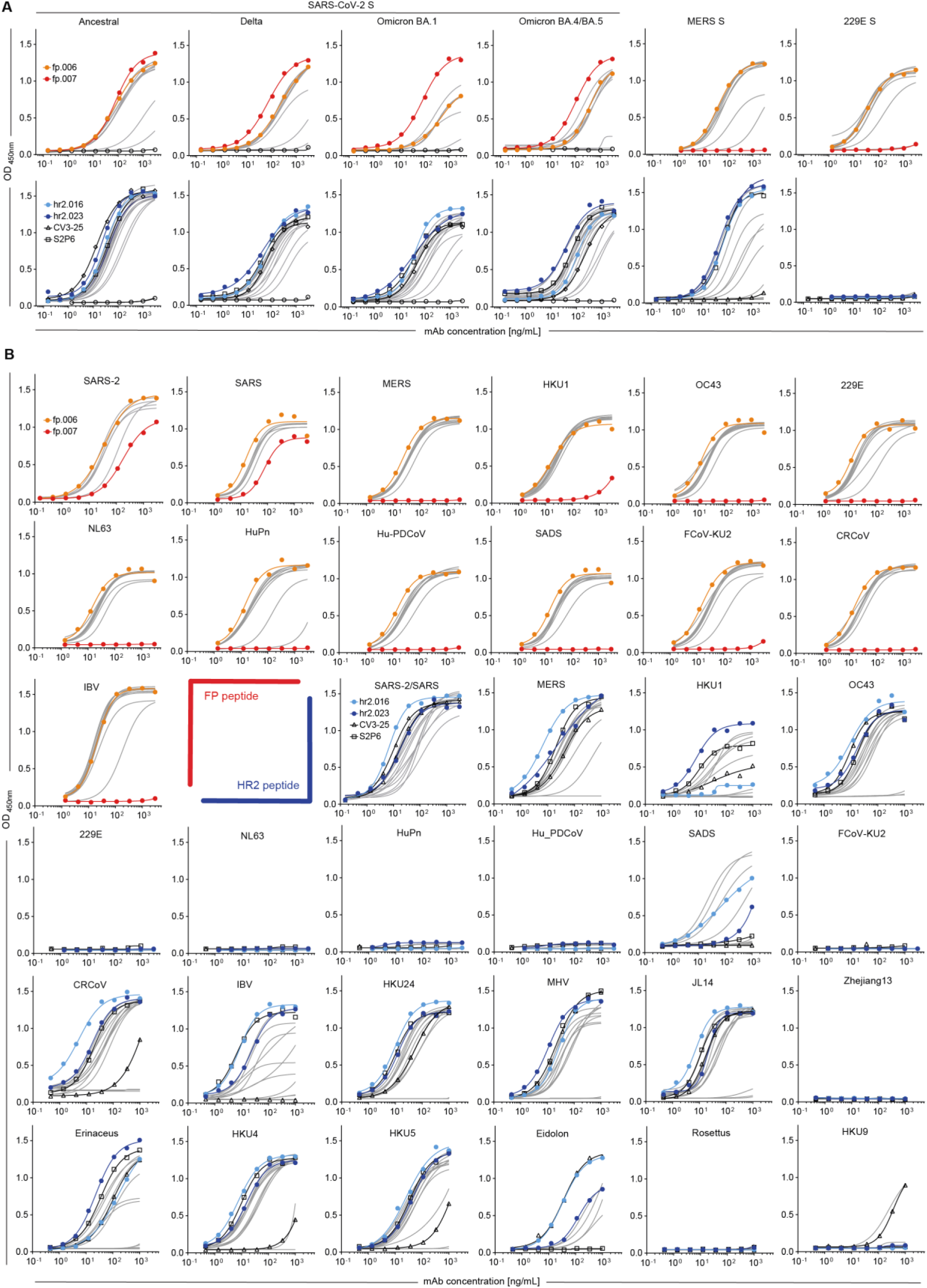
Broadly crossreactive binding of FP and HR2 monoclonal antibodies. (**A** and **B**) Graphs show ELISAs measuring monoclonal antibodies reactivity to S **(A)** and to coldspot peptides **(B)** of coronaviruses. Mean of two independent experiments.

**Fig. S4.**
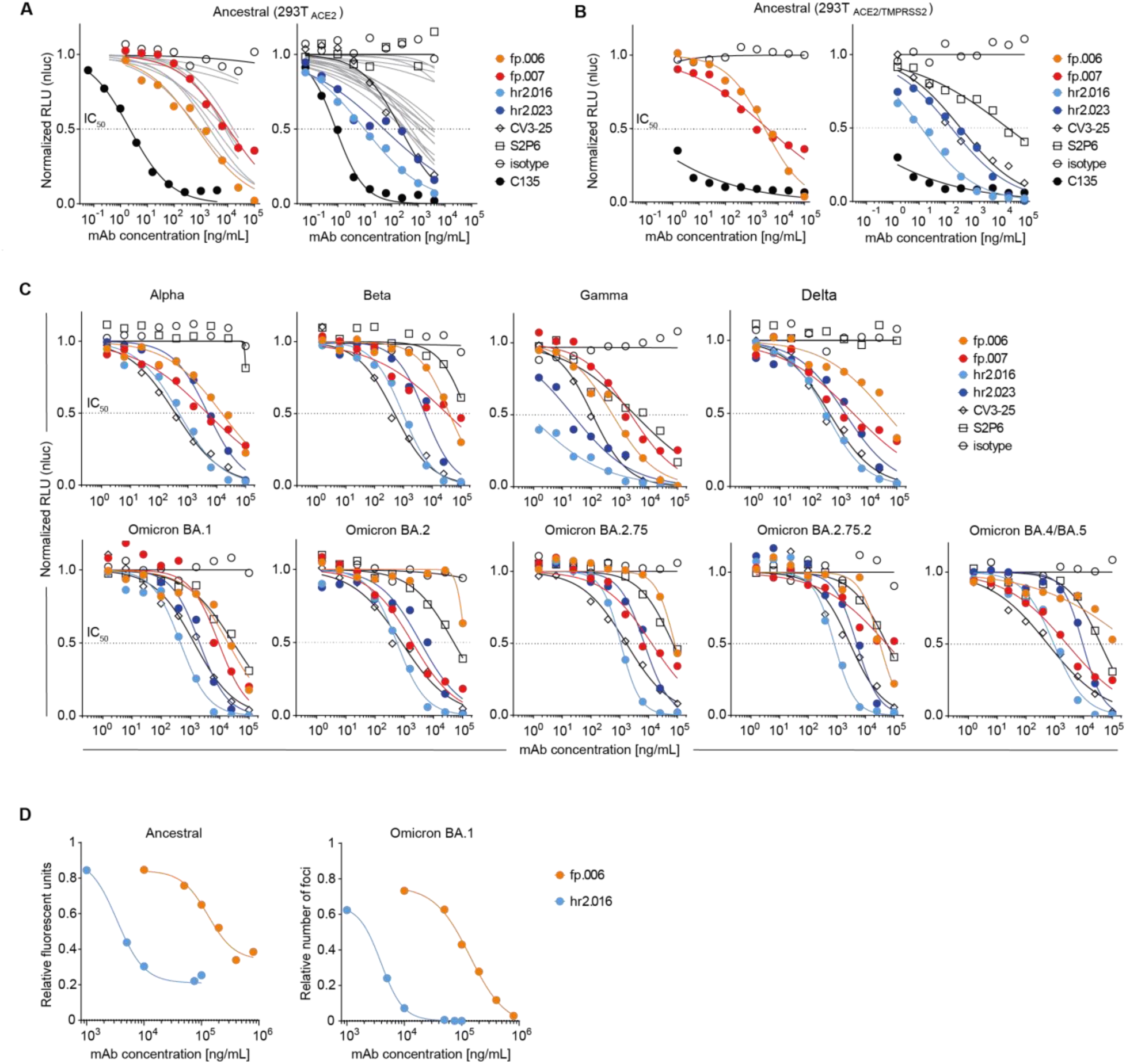
Broad neutralization by FP and HR2 monoclonal antibodies. (**A** and **B**) Neutralization of ancestral SARS-CoV-2 pseudovirus. Graphs show normalized relative luminescence values in cell lysates of 293T_ACE2_ **(A)** or 293T_ACE2/TMPRSS2_ **(B)** cells 48 hours after infection with nanoluc-expressing SARS-CoV-2 pseudovirus in the presence of increasing concentrations of monoclonal antibodies to the FP (top) and HR2 (bottom). Isotype control and antibodies C135 (4), CV3-25 and S2P6 were assayed alongside for comparison; additional antibodies in grey (table S2). Mean of at least two independent experiments. **(C)** Same as in **(A)**, but for SARS-CoV-2 pseudoviruses corresponding to VOC. **(D)** Neutralization of SARS-CoV-2 ancestral and Omicron BA.1 by Focus Reduction Neutralization Test (FRNT).

**Fig. S5.**
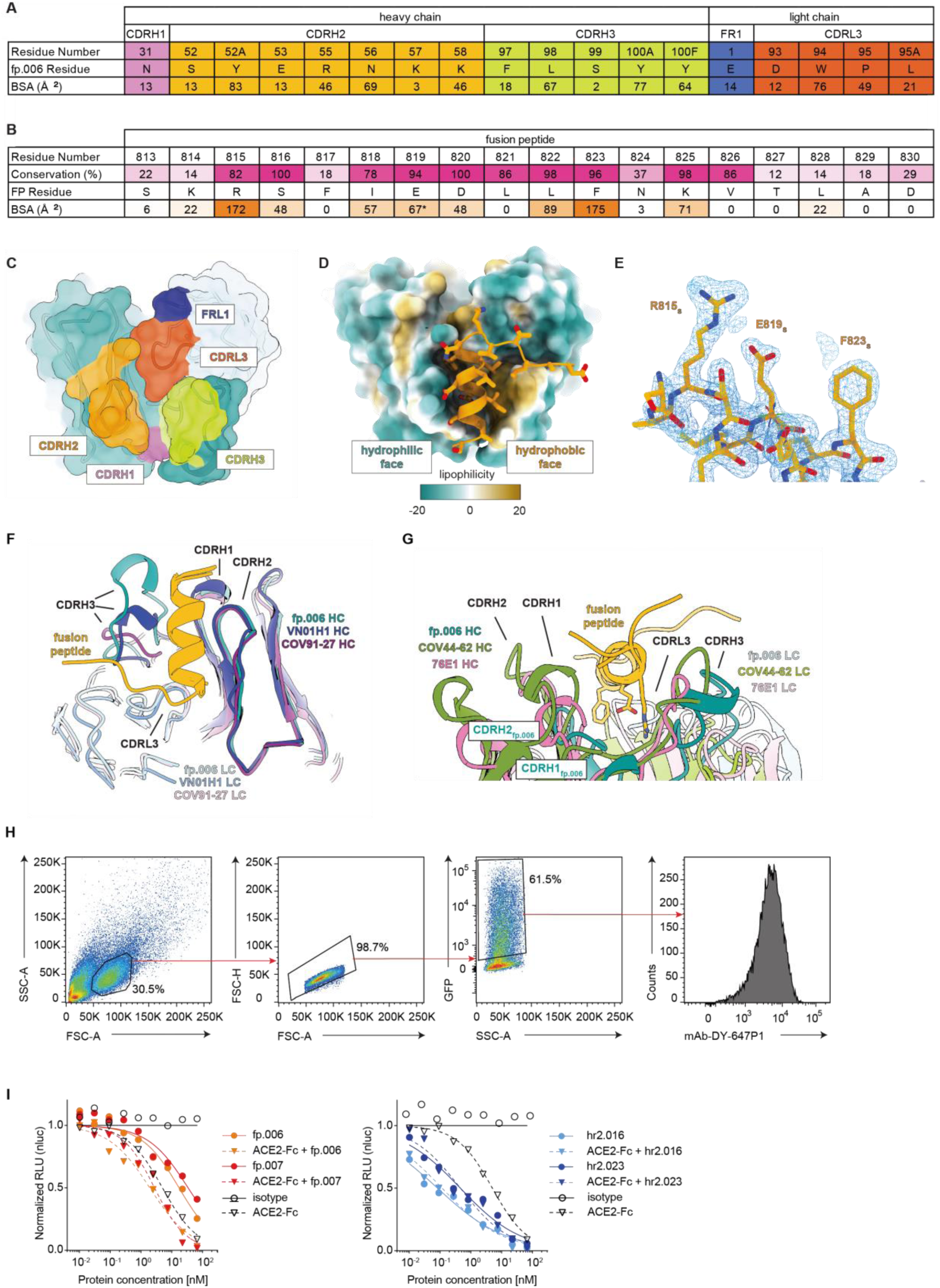
Structure of fp.006 in complex with peptide antigen and effects of ACE2 engagement. **(A)** Contributions of fp.006 heavy and light chain paratope residues to the binding of the fusion peptide. Residue number, identity, and buried surface area (BSA) are shown, and colored as in panel **(C)**. **(B)** Per residue contributions of the FP to the fp.006 epitope. Conservation calculated from the 49 CoV sequences listed in fig. S2B and BSA calculations per FP residue are listed. *Indicates that the BSA calculation does not include contacts mediated by water molecules. **(C)** Surface representation of the fp.006 paratope with interacting residues colored according to their CDR loop identity. Non-interacting heavy chain and light chain residues are colored teal and light teal, respectively. **(D)** Surface representation of the fp.006 paratope with per residue lipophilicity calculations shown. More hydrophobic patches are colored orange and more hydrophilic patches are colored teal, as calculated by ChimeraX. **(E)** Representative electron density for a portion of the FP contoured at 1.7σ. **(F)** Overlay of complex structures between the SARS-CoV-2 fusion peptide and fp.006 (this study; PDB 8D47), VN01H1 (PDB 7SKZ), and COV91-27 (PDB 8D6Z) illustrates a convergent binding mode independent of V gene usage. Structures are aligned on their fusion peptide chains and colored as shown. **(G)** Overlay of complex structures between the SARS-CoV-2 fusion peptide and fp.006 (this study; PDB 8D47), COV44-62 (PDB 8D36), and 76E1 (PDB 7X9E) Fabs, illustrating that anti-FP mAbs of different poses bind to the same face of the amphipathic fusion peptide α-helix, which includes highly-conserved residues, which are shown as sticks. **(H)** Gating strategy for measuring the binding of monoclonal antibodies to the SARS-CoV-2 S expressed by 293T cells by flow cytometry. **(I)** Normalized relative luminescence values in cell lysates of 293T_ACE2_ cells after infection with ancestral SARS-CoV-2 pseudovirus in the presence of increasing concentrations of FP or HR2 antibodies alone or as a cocktail with soluble ACE2. Isotype control in black. For the combination of fp.006+ACE2, P=0.0017 over fp.006 and P=0.0047 over ACE2; for the combination of fp.007+ACE2, P=0.0072 over fp.007 and P=0.2 over ACE2 (n=4; Welch’s t-test, two-tailed).

**Fig. S6.**
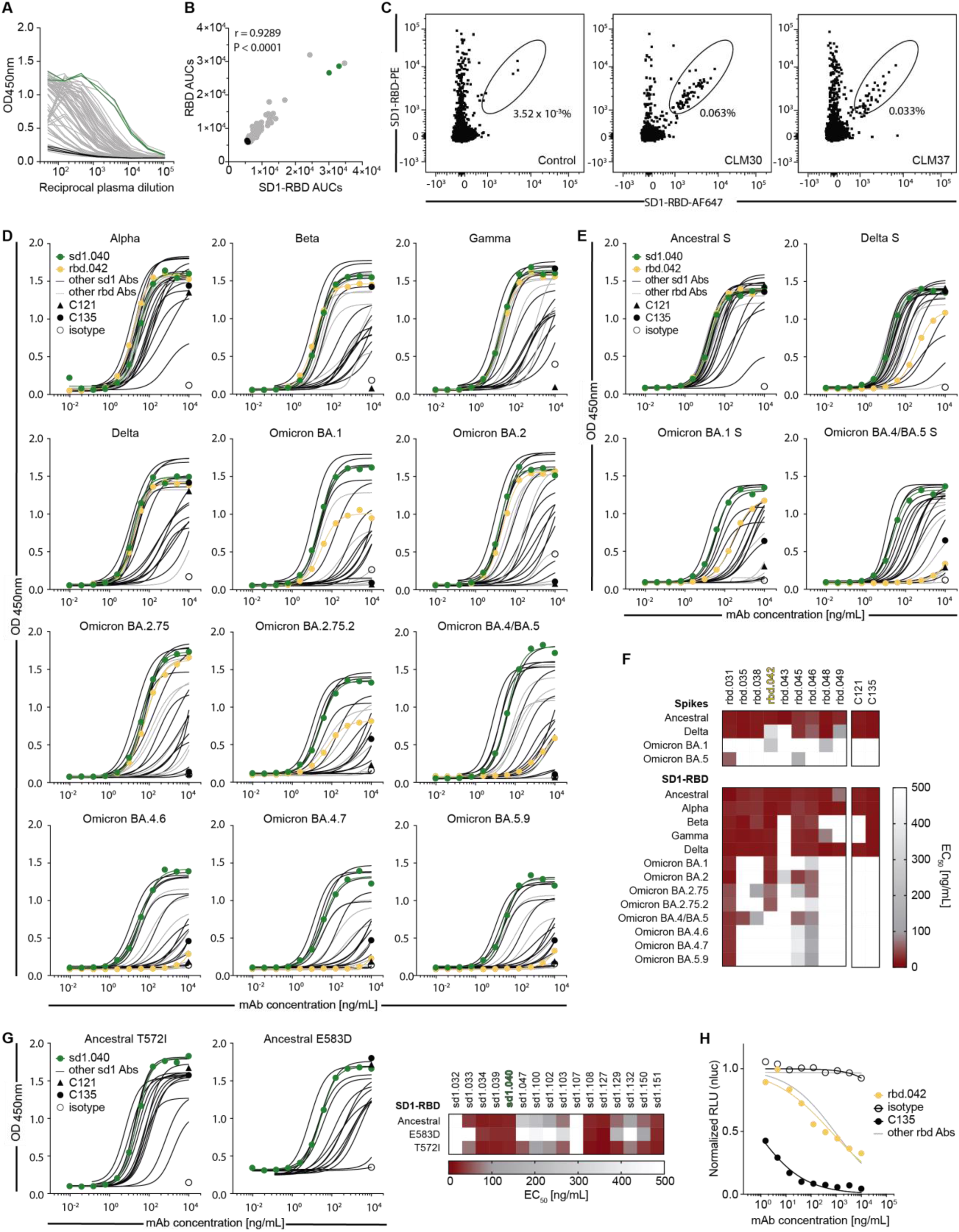
Discovery of SD1 and RBD antibodies. **(A)** Graph shows ELISAs measuring plasma IgG reactivity to RBD. Negative controls in black. Mean of two independent experiments. **(B)** Correlation of plasma IgG reactivities to SD1-RBD and RBD. Pearson correlation. **(C)** Identification of SD1-enriched B cells by flow cytometry. **(D** and **E)** Graphs show ELISAs measuring monoclonal antibodies reactivity to SD1-RBD **(D)** and S **(E)** corresponding to SARS-CoV-2 VOC. Mean of two independent experiments. **(F)** Heatmaps with the binding (EC_50_) of RBD monoclonal antibodies to S (top) or SD1-RBD (bottom) proteins corresponding to SARS-CoV-2 VOC. Two experiments. **(G)** Graphs and summary heatmap for SD1 antibodies binding to SD1-RBD mutant proteins T572I and E583D. **(H)** Neutralization of ancestral SARS-CoV-2 pseudovirus by antibodies to the RBD that are broadly crossreactive to VOC.

**Fig. S7.**
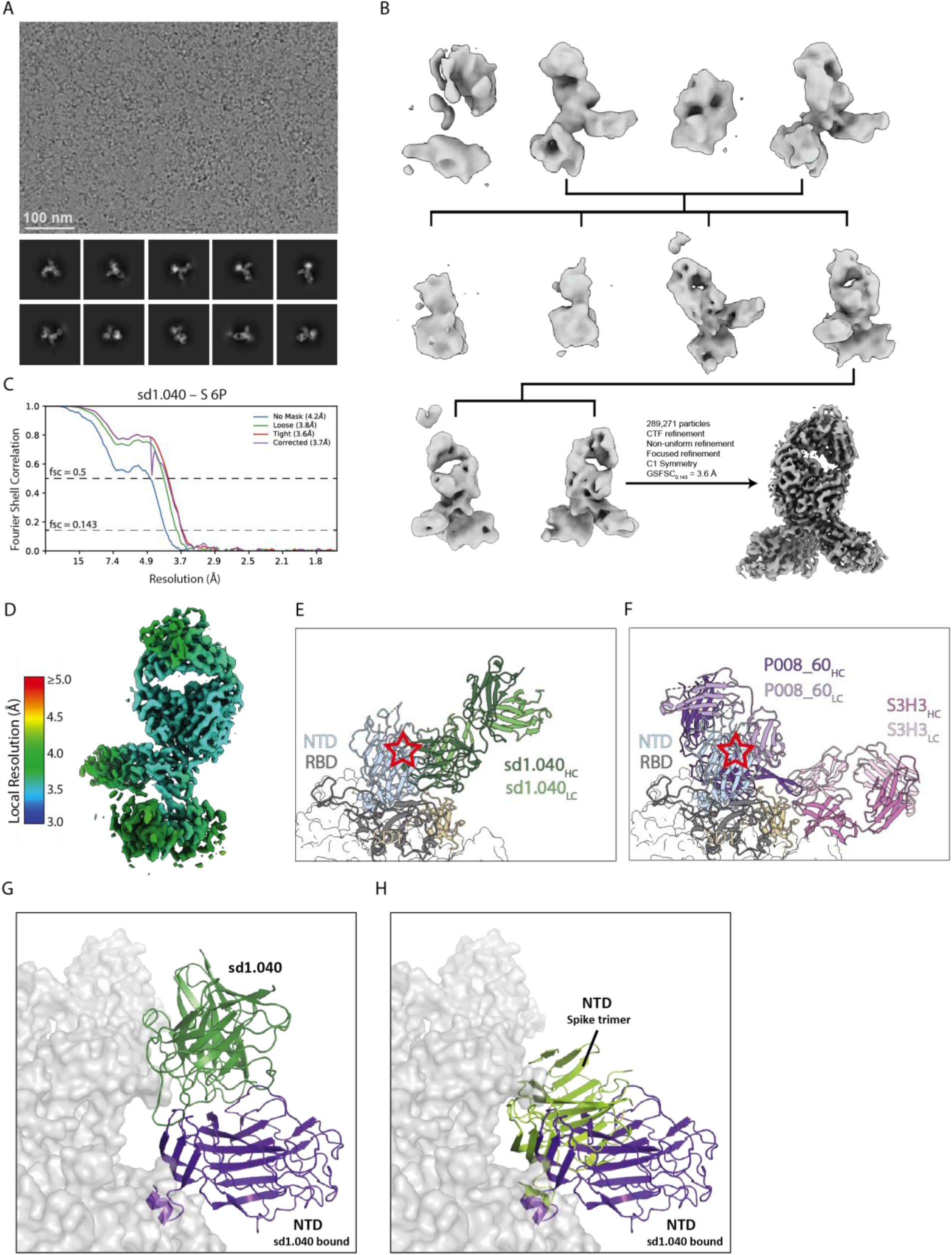
Structure of sd1.040 in complex with S. **(A)** Representative micrograph and 2D class averages selected from the sd1.040-S dataset. **(B)** 3D classification workflow and final refinement strategy. **(C)** Gold-standard FSC plot for the final cryo-EM density map. **(D)** Local resolution estimates calculated in cryoSPARC v3.1. **(E)** Close-up view of the sd1.040 Fab – S complex superimposed onto a prefusion S trimer (PDB 6VXX). Potential clashes with the NTD (cyan) on the neighboring S1 protomer are denoted by a red star. **(F)** Close-up views of the P008_60 Fab (PDB 7ZBU) and S3H3 Fab (PDB 7WD9) superposed onto a prefusion trimeric Spike. Antibody P008_60 shows prominent clashes with the adjacent NTD (red star), while the binding orientation of S3H3 does not clash with neighboring protomers. **(G and H)** According to docking simulations, sd1.040 binds also to the NTD of the adjacent protomer. In **(G)**, NTD moves from the position observed in the experimental structure of the free S to make room for antibody binding (sd1.040 in dark green bound to NTD in purple). In **(H)**, comparison between NTD bound to sd1.040 in simulations (purple) and NTD in the context of free spike trimer (lime).

**Fig. S8.**
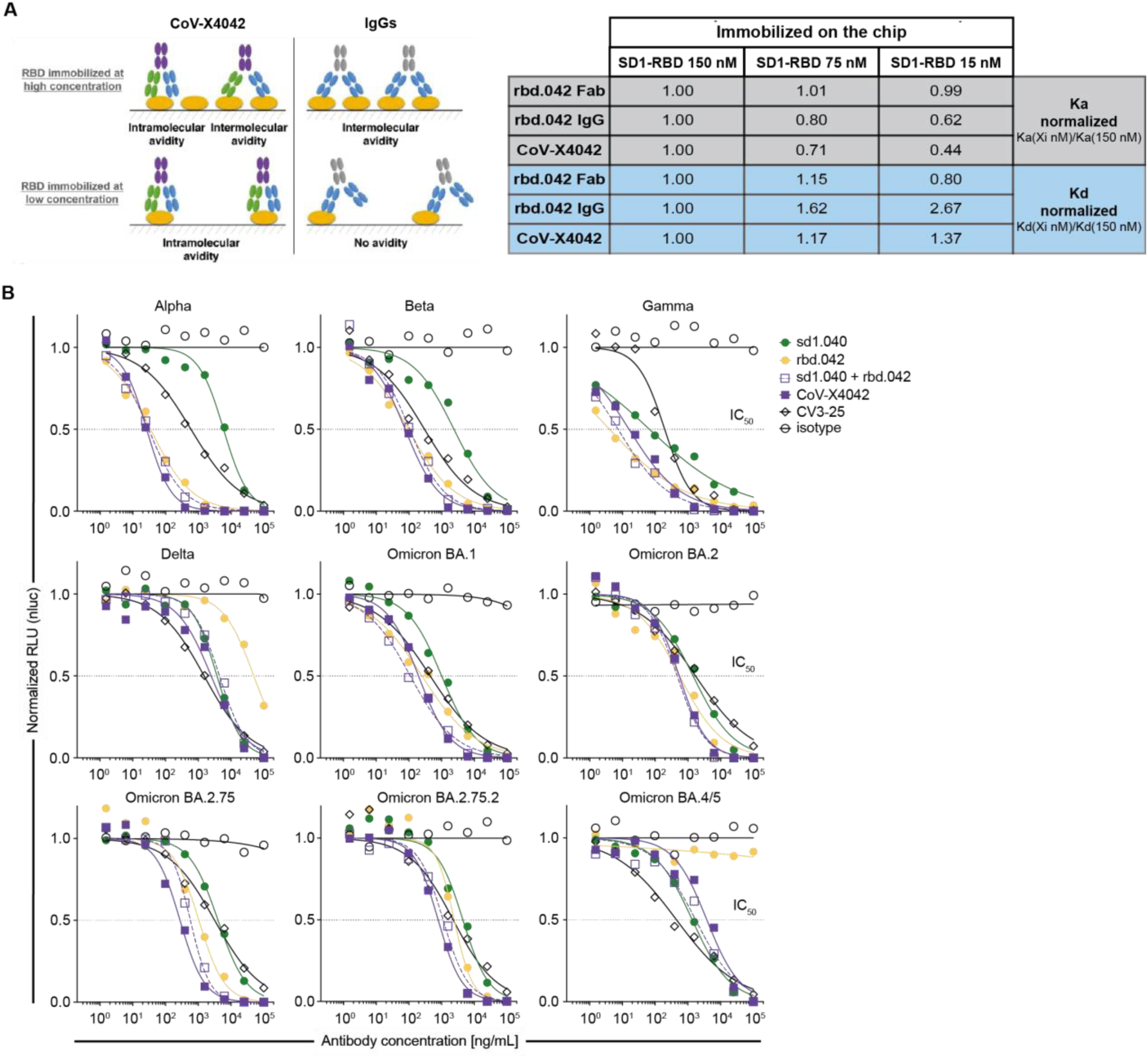
Binding and neutralizing properties of CoV-X4042. **(A)** Both arms of CoV-X4042 bind simultaneously to the same SD1-RBD molecule. Left, diagram showing that at high SD1-RBD concentrations, monoclonal antibodies have avidity effects owing to intermolecular binding (which results in a slower kd), but this is not the case at low SD1-RBD concentrations, because bivalent binding to a single SD1-RBD is impossible. By contrast, the bispecific antibody has avidity at both high and low concentrations, since bivalent binding to its two epitopes on a single SD1-RBD is possible. ka is not affected by avidity. Right, table summarizing the SPR results used to determine ka and kd of IgG, Fab and the bispecific antibody at different concentrations of immobilized SD1-RBD, which confirm experimentally that CoV-X4042 engages bivalently on a single SD1-RBD. **(B)** Neutralization of VOC by CoV-X4042 and parental monoclonal antibodies.

